# Multiomic screening platform uncovers the impact of histone mutations on chromatin and cell fate

**DOI:** 10.64898/2026.04.14.718582

**Authors:** Ziyang Ye, Alireza Khademi, Renée L. Barbosa, Foster Birnbaum, Margaret R. Brown, Colin E. Fowler, Arianna Arroyo-Ortega, Daniel Lee, Lijuan Feng, Leah A. Gates, Agata L. Patriotis, Amy E. Keating, Francisco J. Sánchez-Rivera, Yadira M. Soto-Feliciano

## Abstract

Somatic missense mutations in histone genes, often referred to as ‘oncohistones’, have been identified in diverse types of human cancers. The functional and mechanistic impact of most oncohistones remains unknown. To address this gap, we developed CHANCLA, a modular platform for high-throughput functional screening of oncohistones using multiomic phenotypic readouts. We used CHANCLA to systematically measure the impact of 303 human oncohistones on cellular proliferation, differentiation, histone-specific post-translational modifications, and chromatin accessibility. Integrative multiomic analyses revealed discrete oncohistone molecular classes that promote proliferation, block lineage-specific differentiation, and physically remodel the chromatin landscape by altering specific histone modifications and reducing nucleosome stability. Structural mapping and computational modeling studies uncovered that functionally convergent mutations are clustered at key nucleosome interfaces, particularly H2B-H4, and that chromatin accessibility-promoting mutations are linked to mono-nucleosome destabilization. Leveraging this multiomic resource, we discovered that the H3.3-Q5H mutant histone is a *bona fide* human oncohistone that accelerates lung adenocarcinoma growth *in vivo*. Mechanistically, we found that *H3*.*3-Q5H* expression leads to suppression of promoter-associated H3K4me3 and expansion of repressive H3K27me3 domains, resulting in increased KRAS signaling and gene expression programs associated with epithelial-to-mesenchymal transition. Together, this work provides a multiomic functional atlas of cancer-associated histone mutations, identifies structural and mechanistic principles governing chromatin reprogramming by oncohistones, and establishes CHANCLA as a modular platform for systematic discovery of mechanisms and vulnerabilities associated with these genetic lesions.

## INTRODUCTION

The histone octamer organizes eukaryotic genomes into chromatin, and its post-translational modification is a central mechanism of gene regulation.^1^ Disruption of chromatin regulatory systems, including DNA methyltransferases, histone-modifying enzymes, chromatin remodelers, and PTM readers, is a hallmark of human cancers.^2–6^ Beyond alterations in chromatin regulators, somatic mutations in genes encoding histones have recently emerged as a direct mechanistic contributor to chromatin misregulation in cancer.^7^ Termed ‘oncohistones’, these mutations were first identified in pediatric gliomas and bone tumors, where highly recurrent lysine-to-methionine (K→M) substitutions in histone H3 (*H3K27M* and *H3K36M*) act as dominant drivers by globally suppressing their cognate histone modification in *trans*.^*8–10*^ These discoveries established that single amino acid changes in histones can reprogram chromatin states and cell fate, with profound consequences for tumorigenesis.^11–13^

Large-scale sequencing efforts have since revealed that histone mutations extend far beyond the H3 K→M paradigm. Hundreds of somatic missense mutations have been catalogued across genes encoding all four core histones and the H3.3 variant in diverse adult and pediatric malignancies.^14,15^ Structural mapping studies have placed these mutations and their respective wild-type residues at histone-DNA contacts, histone-histone interfaces, and PTM sites throughout the nucleosome ^14–16^. However, mechanistic experimental studies of mammalian oncohistones, including efforts to systematically investigate their functional impact on human cell physiology, have lagged behind. In fact, most cancer-associated histone mutations have never been experimentally investigated to assess their effects on chromatin regulation, cellular fitness, or lineage commitment. The gap between genomic annotation and functional mechanistic understanding represents the main barrier to determining which histone mutations are oncogenic drivers in human cancer and how they lead to malignant chromatin dysregulation.

Here, we address this gap by developing CHANCLA (<u>C</u>ancer-associated <u>H</u>istone <u>A</u>lteration <u>N</u>etworks and <u>C</u>ellular <u>L</u>andscape <u>A</u>nalysis), a modular and scalable platform that integrates flexible barcoded gene expression constructs with multiomic functional profiling for multidimensional high-throughput phenotyping of oncohistones. As a proof-of-concept, we constructed a CHANCLA library to systematically measure the functional impact of 303 human oncohistones on cellular proliferation, differentiation, histone-specific post-translational modifications, and chromatin accessibility (CHANCLA-303). Integrative multiomic analyses revealed discrete oncohistone molecular classes that promote cellular proliferation, block lineage differentiation, and remodel the chromatin landscape by altering specific histone modifications and reducing nucleosome stability. Structural mapping and computational analyses showed that functionally convergent mutations are clustered at key nucleosome interfaces, particularly the H2B-H4 interface, and that mutations promoting mono-nucleosome destabilization lead to increased chromatin accessibility. Leveraging this multiomic resource, we identified and validated the *H3*.*3-Q5H* mutant histone as a *bona fide* human oncohistone that accelerates lung adenocarcinoma growth *in vivo*. Mechanistically, we found that *H3*.*3-Q5H* expression leads to suppression of promoter-associated H3K4me3 and expansion of repressive H3K27me3 domains, which in turn lead to increased levels of KRAS signaling and gene expression programs associated with epithelial-to-mesenchymal transition. Together, these studies define a multiomic functional landscape of human cancer-associated histone mutations, reveal structural and mechanistic principles by which oncohistones reprogram mammalian chromatin and cellular behavior, and establish a scalable platform for systematic identification of mechanisms and vulnerabilities associated with oncohistones.

## RESULTS

### Pan-cancer landscape of somatic histone mutations

To define the spectrum of somatic histone mutations in human cancer, we analyzed 11,685 tumors from the MSK-IMPACT and TCGA cohorts, focusing on 40 genes encoding canonical H2A, H2B, H3, and H4, as well as variant H3.3 histones (**Figure 1A**). These genes are primarily clustered on chromosomes 1, 6, and 17 and include multiple paralogs within each family. To identify recurrent mutations likely to be under positive selection, we filtered for missense variants present in ≥ 3 patients. This analysis yielded 303 distinct recurrent missense mutations distributed across all five histone families (**Figure 1B**).

**Figure 1:**
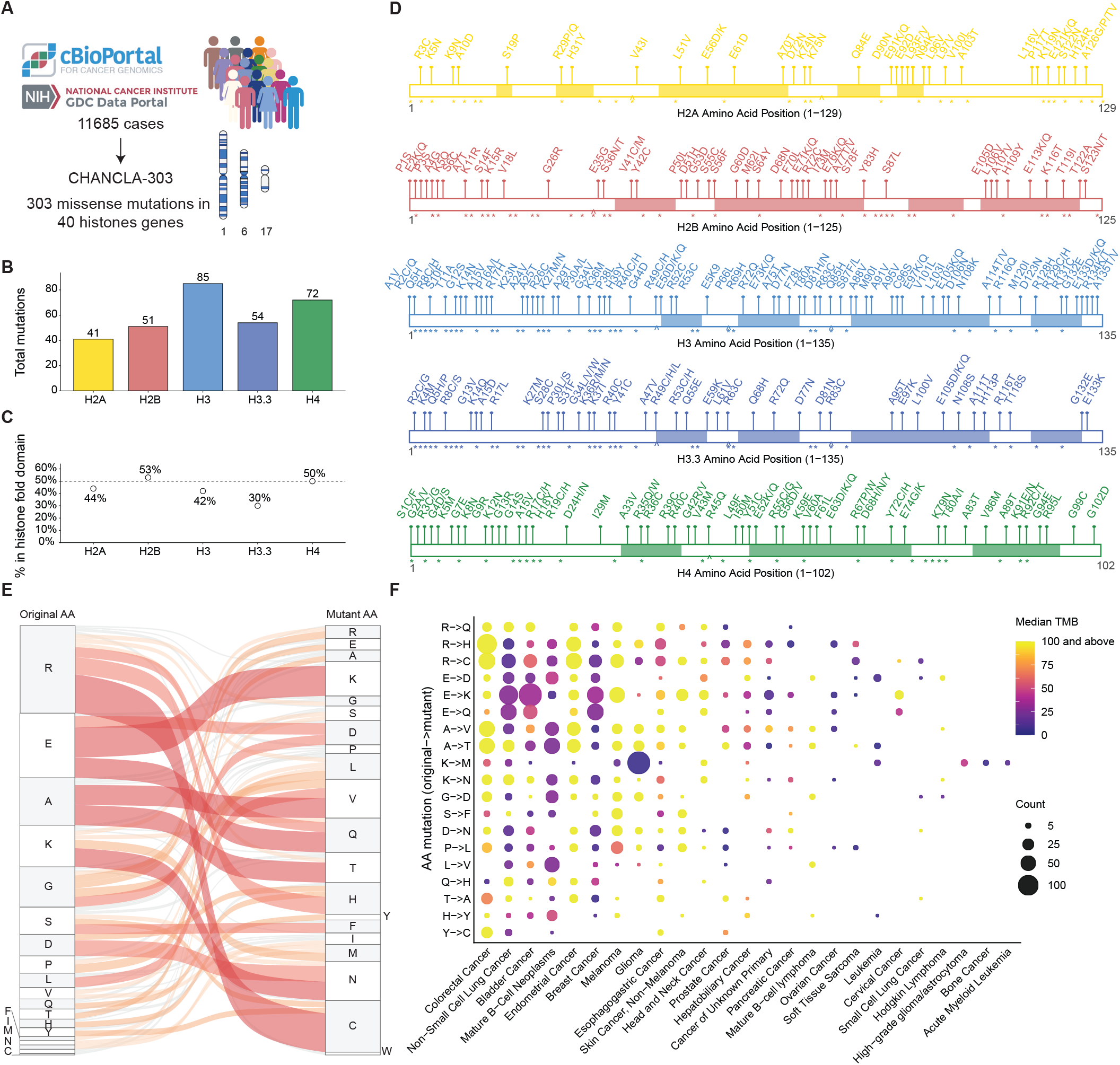
Landscape of cancer-associated histone mutations and design of CHANCLA-303. **(A)** Schematic of oncohistone cDNA library generation. **(B)** Distribution of mutations in the library across histone proteins. **(C)** Distribution of mutations in the library across histone fold domains. **(D)** Visualization of mutations in the library across their respective histone proteins. Shaded area indicates histone fold domains. (*) indicates residues reported to have PTMs. (^) indicates residues contacting nucleosomal DNA. **(E)** Sankey plot of amino acid mutations (original AA → mutant AA) in the oncohistone library. **(F)** Major cancer type distributions for top enriched amino acid mutation patterns. Dot size represents patient counts, colored by sample median tumor mutational burden (TMB).

Mutation burden was unevenly distributed. H3-encoding genes harbored the largest fraction of total mutations (85 in H3 and 54 in H3.3), followed by H4 (72), H2B (51), and H2A (41) (**Figure 1B**). Mapping these substitutions onto linear protein sequences revealed that mutations spanned both intrinsically disordered N-terminal tails and structured histone-fold domains (**Figure 1C-D**). However, the proportion of mutations within the histone fold differed markedly by family: H2B (53%) and H4 (50%) carried roughly half their mutations in the globular domain, whereas H3 (42%), H2A (44%), and H3.3 (30%) were skewed toward the tails (**Figure 1C**). These inter-family differences indicate that distinct structural compartments of the nucleosome are selectively targeted depending on histone identity.

We next examined the spectrum of amino acid substitutions to assess whether specific biochemical changes were overrepresented (**Figure 1E**). Substitution classes were non-uniform across the dataset. Arginine (R) substitutions were the most frequent, predominantly R→cysteine (C), R→histidine (H), or R→glutamine (Q). Given the central roles of arginine residues in histone-DNA contacts,^17^ nucleosome stability,^18^ and post-translational modifications,^19^ this enrichment is consistent with selective targeting of functionally constrained positions rather than neutral mutational drift. Glutamic acid-to-lysine (E→K) substitutions, which introduce charge reversal, and lysine-to-methionine (K→M) substitutions, which eliminate methylation potential, were also recurrent. The predominance of changes that alter residue charge, hydrogen-bonding capacity, or PTM competence suggests that many cancer-associated histone mutations are selected for their ability to disrupt chromatin structure and regulatory interfaces.

To determine whether paralogous histone genes encoding nearly identical proteins share or diverge in their hotspot distributions, we mapped mutation frequencies along each paralog’s amino acid sequence (**Supplementary Figure 1**). Despite high sequence conservation, hotspot residues were frequently non-overlapping across paralogs. Within the H2A family, R35 mutations were concentrated in *H2AC1*, whereas A126 mutations predominated in *H2AC17* (**Supplementary Figure 1A**). In the H2B family, E76 emerged as a dominant hotspot in *H2BC4* and *H2BC5* but was not comparably enriched in other H2B paralogs (**Supplementary Figure 1B**). Among canonical H3.1 genes, E97 was preferentially enriched in *H3C1, H3C3, H3C4, H3C6, H3C8, H3C10, H3C11, and H3C12*, while E105 was enriched in *H3C2, H3C3, H3C4, H3C6, and H3C7*; other H3 paralogs harbored distinct hotspots (**Supplementary Figure 1C**). The two H3.3-encoding genes were particularly divergent: *H3-3A* was strongly enriched for K27 and G34 substitutions, consistent with prior reports in pediatric glioma,^20,21^ whereas *H3-3B* accumulated mutations at alternative sites (**Supplementary Figure 1C**). H4 paralogs similarly displayed gene-specific hotspot distributions (**Supplementary Figure 1D**). These patterns indicate that mutation selection operates not only at the residue level but also at the level of individual histone genes, even when encoded proteins are nearly identical.

We next assessed whether specific histone genes were preferentially mutated in particular cancer types and whether these events occurred in distinct tumor mutational burden (TMB) contexts. For each gene, we quantified mutation frequency across tumor types and overlaid median sample TMB (**Figure 1F**; **Supplementary Figure 2**). Consistent with prior studies, K→M substitutions in *H3-3A* were enriched in low-TMB gliomas, reflecting the canonical *H3K27M* driver mutation.^20,21^ Beyond this established example, gene- and cancer-type-specific biases were widespread. Within the H2A family, *H2AC1* mutations were enriched in melanoma, whereas *H2AC6, H2AC11, H2AC15, H2AC16*, and *H2AC17* were more frequently mutated in mature B-cell neoplasms with relatively low TMB (**Supplementary Figure 2A**). Among H2B paralogs, *H2BC5* showed strong enrichment in colorectal, non-small cell lung, and bladder cancers (**Supplementary Figure 2B**).

While H3 family mutations were associated with lower median TMB, H2A, H2B, and H4 mutations occurred more frequently in tumor types with higher background mutational burdens (**Supplementary Figure 2C**). This divergence suggests that H3 alterations are more often selected in genomically quieter tumors, consistent with stronger positive selection in less rapidly evolving contexts. Within H3.1 genes, *H3C2, H3C4, H3C8*, and *H3C12* mutations were enriched in melanoma, while *H3C8* mutations also showed enrichment in mature B-cell neoplasms (**Supplementary Figure 2C**). *H3-3A* mutations were strongly associated with glioma, whereas *H3-3B* exhibited a distribution more closely resembling canonical H3 genes (**Supplementary Figure 2C**). Because comprehensive sequencing of H4 genes has only recently been incorporated into clinical panels, H4 mutation patterns may be underpowered for detecting additional cancer-type associations (**Supplementary Figure 2D**).

Together, these analyses reveal multiple layers of non-random structure in the somatic histone mutation landscape. Biases are evident at the level of substitution class, structural domain, residue-specific hotspot, paralog identity, and tumor-type enrichment. Despite high sequence conservation, histone mutations are neither interchangeable across paralogs nor uniformly distributed across cancers; instead, they reflect gene-specific evolutionary constraints and cancer-type-specific selective pressures. These observations establish a framework for systematic functional interrogation of oncohistones across cellular and chromatin contexts.

### Pooled screening identifies histone mutations that promote proliferation and impair differentiation

Several histone mutations have been shown to influence cellular fitness and lineage plasticity in specific tumor contexts.^8,9,22^ However, the functional consequences of most cancer-associated histone mutations remain unknown, particularly beyond well-characterized K→M substitutions in histone H3.^7^ To systematically interrogate this, we generated a pooled lentiviral cDNA library encoding all 303 recurrent missense mutations identified in our pan-cancer analysis (**Figure 1**). Each mutation was encoded within its native human histone gene sequence and linked to a unique 15-bp DNA barcode for sequencing-based deconvolution. Wild-type H2A, H2B, H3, H3.3, and H4 constructs were included as internal controls. Because canonical histone mRNAs lack polyadenylation (pA) tails,^23^ we appended a pA signal to enable transcript capture and cloned the cDNAs in antisense orientation relative to the viral backbone to prevent transcriptional interference^24^ (**Supplementary Figure 3A**). This barcoded oncohistone library, CHANCLA-303, provides a scalable, internally controlled platform for high-throughput functional analysis of cancer-associated histone mutations.

#### Proliferation screening

To assess how oncohistone expression affects cellular fitness, we performed pooled proliferation screens in C3H10T1/2 mouse mesenchymal stem cells (MSCs), an established model system for oncohistone studies.^9,10^ Cells were transduced with CHANCLA-303 at low multiplicity of infection (MOI ∼0.3), and barcode abundance was tracked across twelve serial passages (**Figure 2A**; **Supplementary Figure 3A**). For each construct, normalized abundance was converted to passage-wide z-scores, and a linear regression slope across all passages was calculated to quantify relative proliferative enrichment or depletion (**Figure 2B**). Significant hits (see **Methods**) were identified across all histone families: 13 in H2A, 34 in H2B, 34 in H3, 23 in H3.3, and 17 in H4 (**Figure 2B**). Individual validation experiments confirmed that selected hits recapitulated the direction and magnitude of pooled screen slopes (**Supplementary Figure 3B**), demonstrating the quantitative reliability of this systematic approach.

**Figure 2:**
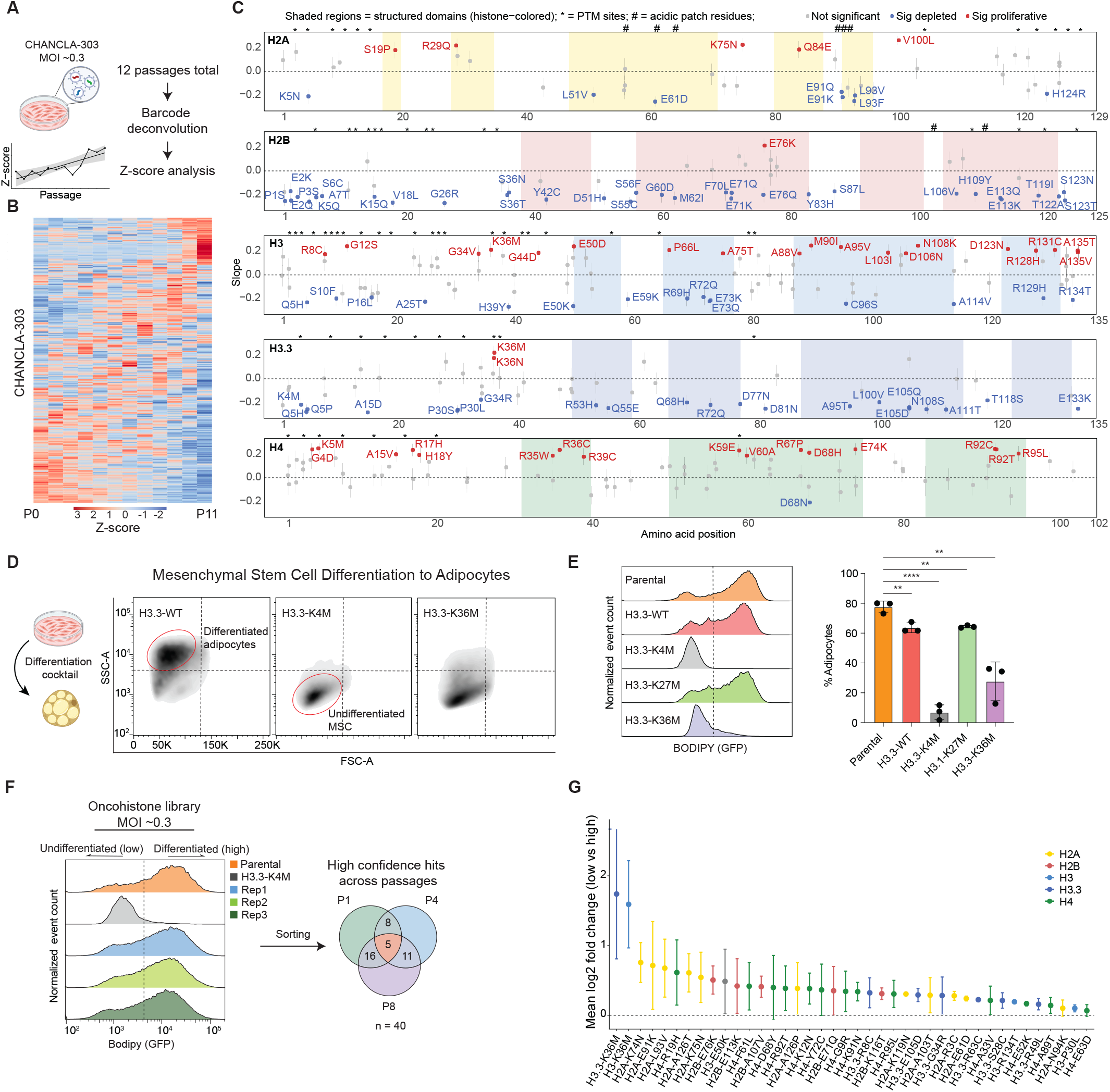
High-throughput functional screening identifies oncohistone mutations driving proliferation and differentiation blockade. **(A)** Schematic of oncohistone proliferation screen and analysis. **(B)** Heatmap shows normalized Z-score for each oncohistone (per row) across Passage 0 to 11. **(C)** Plot of the fitted slope of each oncohistone’s normalized Z-score (*P*-value < 0.05) along the amino acid position of histone proteins. Shaded area indicates histone fold domains. (*) indicates residues reported to have PTMs. (#) indicates acidic patch residues. **(D)** Representative flow cytometry density plots of MSCs subjected to adipocyte differentiation cocktail. **(E)** Normalized event count of MSCs subjected to adipocyte differentiation cocktail and stained by BODIPY. Percentage of differentiated adipocytes gated based on *H3*.*3-K4M*-MSCs BODIPY signal. **(F)** Schematic of oncohistone differentiation screen and differentiation blocker hits identification process. **(G)** Mean log_2_(fold-change) of differentiation blocker hits from the screen, colored by histone family.

Proliferation phenotypes were highly residue- and domain-specific (**Figure 2C**). In H2A, mutations in the αN domain, α1 helix, and α3 helix (e.g., *S19P, R29Q, Q84E*) consistently conferred proliferative advantage, whereas substitutions in the α2 helix and αC domain were strongly depleted. Mutations affecting H2A acidic patch residues, which mediate internucleosomal interactions and higher-order chromatin organization,^25,26^ were uniformly deleterious, underscoring the essential structural role of this surface. Most H2B mutations were similarly poorly tolerated, indicating strong functional constraints within this histone. However, *H2B-E76K* was a clear outlier and one of the strongest proliferative hits across the entire library, consistent with its recurrent occurrence in bladder and head-and-neck cancers and its previously reported growth-promoting effects in mammary epithelial cells.^22^ Unlike H2A acidic patch substitutions, E76 lies at the H2B-H4 dimer-tetramer interface,^17^ and its mutation likely destabilizes nucleosome architecture without abolishing essential chromatin-binding contacts. The selective enrichment of *H2B-E76K* therefore suggests that disruption of specific histone-histone interfaces, rather than global nucleosome destabilization, can create a narrow mutational window permissive for enhanced proliferation of cancer cells.

H3 and H3.3 mutations produced the most heterogeneous proliferation phenotypes. Substitutions at H3A135 (*A135T/V*), enriched in mature B-cell neoplasms, conferred robust proliferative advantage (**Figure 2C**; **Supplementary Figure 2C**). As expected, *H3-K36M* and *H3*.*3-K36M* were strong proliferative enrichers, consistent with their established roles in undifferentiated sarcoma and chondroblastoma.^9,13^ Conversely, *H3*.*3-K4M* was strongly depleted, matching previous reports of its anti-proliferative effects in preadipocyte models.^27^ Outside these defined hotspots, most H3.3 mutations were poorly tolerated in MSCs, suggesting that the H3.3 variant is less permissive of random perturbation in this cellular context (**Figure 2C**). Unexpectedly, a substantial fraction of H4 mutations promoted proliferation. *H4-R92C* and *H4-R92T* ranked among the strongest enrichers across the entire screen (**Figure 2C**). H4R92 forms stabilizing hydrogen bonds with H2BE76,^17^ and disruption of either residue yielded comparable proliferation phenotypes, indicating that perturbation of a shared structural interface generates convergent functional outcomes.

To examine whether specific biochemical substitution classes were associated with fitness effects, we grouped mutations by amino acid gained or lost and calculated average proliferation slopes (**Supplementary Figure 3C-D**). Loss of arginine (n = 18) and gain of cysteine (n = 8) were disproportionately associated with proliferative enrichment, suggesting that disruption of positively charged DNA-contacting residues or introduction of reactive thiol side chains can confer a growth advantage. To explore the structural basis of these phenotypes, we mapped residues with significant proliferative enrichment onto a pairwise Cα atom distance matrix derived from a high-resolution nucleosome structure^14,28^ (**Supplementary Figure 4**). Enriched residues clustered at spatially proximal positions within and across histone subunits, including multiple sites in the H3 and H4 histone-fold domains and at the H2B-H4 interface (H2BE76, H4D68, H4R92). These spatial relationships suggest that cancer-associated histone mutations can exert growth-promoting effects through coordinated perturbation of key nucleosome-stabilizing contacts rather than through isolated structural changes.

#### Differentiation screening

Mesenchymal stem cell models have been widely used to study how oncohistones influence cell fate decisions, revealing that specific histone mutations can disrupt differentiation programs and maintain progenitor-like states.^9,12,29^ However, conventional assays for mesenchymal differentiation rely on fixation, dye extraction, and bulk readouts,^30^ which limit throughput and preclude single-cell and live-cell analyses. To overcome these limitations, we developed a flow cytometry-based assay that quantifies adipogenic differentiation at single-cell resolution using BODIPY staining of lipid accumulation as a measure of mesenchymal differentiation into the adipocytic lineage^30,31^ (**Figure 2D**). As validation, we confirmed that MSCs expressing *H3*.*3-K4M* or *H3*.*3-K36M* showed marked reductions in adipocyte differentiation relative to parental or wild-type H3.3 controls (**Figure 2D-E**), consistent with previous studies.^12,27^ Reduced expression of the adipogenic transcription factors *Cebpα* and *Pparγ*^*32*^ independently confirmed impaired differentiation (**Supplementary Figure 5A**), further validating the sensitivity of the assay.

We then adapted this system for pooled screening. CHANCLA library-transduced MSCs were subjected to adipogenic induction, and BODIPY-high (differentiated) and BODIPY-low (undifferentiated) populations were sorted at multiple passages (P1, P4, and P8) (**Figure 2F, Supplementary Figure 5B**). Barcode representation was quantified by sequencing, and mutations reproducibly enriched in the undifferentiated fraction across passages were designated as high-confidence differentiation-blocking hits (see **Methods**). This analysis identified 40 mutations distributed across all four core histone families and the H3.3 variant (**Figure 2G**). We individually transduced cDNAs from 15 hits into MSCs, and all were confirmed to block differentiation to varying degrees (**Supplementary Figure 5C**). Among the strongest hits were *H2B-E76K* and *H4-R92T*, mirroring their proliferative phenotypes and reinforcing the functional importance of the H2B-H4 interface. Additional differentiation-blocking mutations included *H2A-K74N, H2A-K75N*, and several H4 histone-fold domain substitutions, indicating that discrete structural perturbations can broadly impair lineage commitment. Overlaying differentiation-blocking hits onto the nucleosome Cα distance matrix revealed spatial clustering at the H2B-H4 interface (**Supplementary Figure 6**), paralleling the pattern observed for proliferation-promoting mutations (**Supplementary Figure 4**).

Together, the pooled proliferation and differentiation screens demonstrate that cancer-associated histone mutations exert highly selective, residue-dependent effects on cellular fitness and lineage commitment. While most mutations were poorly tolerated, a distinct subset including canonical H3K36 substitutions and mutations at the H2B-H4 nucleosome interface consistently promoted proliferation and blocked differentiation. The spatial clustering of these mutations at structurally interacting residues within the nucleosome core suggests that perturbation of nucleosome architecture represents a shared mechanism underlying these convergent phenotypes.

### Chromatin screens identify oncohistones that alter histone modifications and nucleosome stability

Because several mutations that altered proliferation and differentiation clustered at residues involved in chromatin regulation or nucleosome stability, we next asked whether cancer-associated histone mutations broadly reshape chromatin states. To address this, we developed a pooled intracellular staining strategy that quantifies global histone post-translational modifications (PTMs) by flow cytometry, followed by barcode deconvolution to identify mutations associated with altered PTM levels (**Figure 3A**). MSCs expressing the oncohistone library were stained with antibodies recognizing five representative chromatin marks: enhancer-associated H3K4me1, promoter-associated H3K4me3, constitutive heterochromatin-associated H3K9me3, facultative heterochromatin-associated H3K27me3, and transcription-associated H3K36me3. For each mark, cells within the highest and lowest ∼5% of fluorescence intensity were isolated by FACS and sequenced at multiple passages (P1 and P5). Mutations reproducibly enriched across passages were designated as high-confidence hits (**Figure 3A**; see **Methods**).

**Figure 3:**
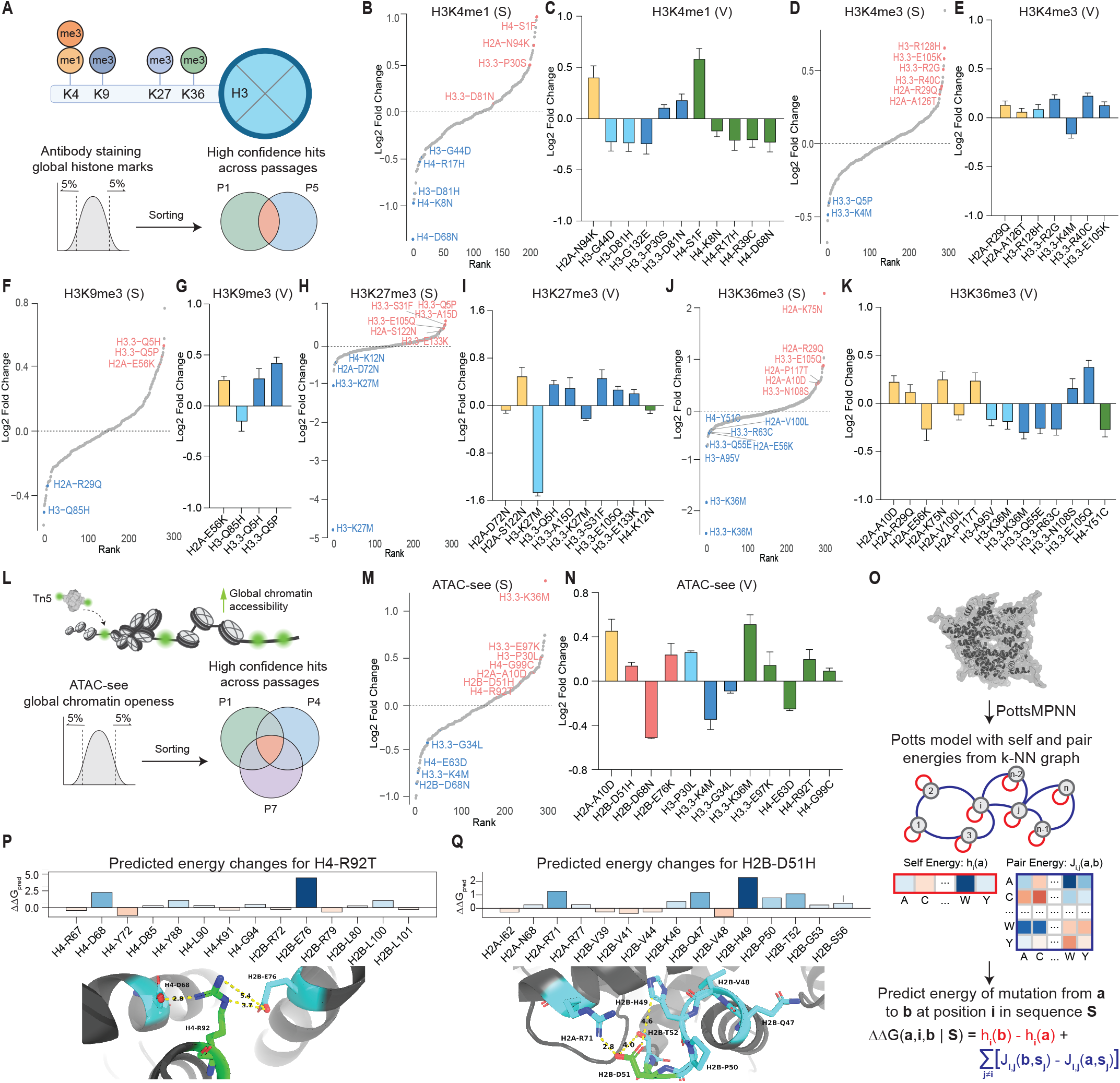
Oncohistone mutations reprogram chromatin states and gene expression. **(A)** Schematic of oncohistone global histone PTM screen and hits identification process. **(B**,**D**,**F**,**H**,**J)** Plots of log_2_(fold-change) (top 5% vs bottom 5%) of oncohistone global histone PTM screen data (H3K4me1, H3K4me3, H3K9me3, H3K27me3, H3K36me3), labeled as S. **(C**,**E**,**G**,**I**,**K)** Plots of log_2_(fold-change) of oncohistone hits ICS validation data (normalized to EV; P-value < 0.05), labeled as V. **(L)** Schematic of oncohistone global chromatin accessibility screen and hits identification process. **(M)** Plots of log_2_(fold-change) of oncohistone global chromatin accessibility screen data. **(N)** Plots of log_2_(fold-change) of oncohistone hits validation data (normalized to EV; *P*-value < 0.05). **(O)** Schematic of how PottsMPNN generates a Potts model containing self and pair energies that can be used to predict the effect of mutations on protein stability. **(P)** PottsMPNN predictions for how pair energies change upon the H4-R92T mutation. Structural analysis of the neighboring residues highlights that interactions with H4D68 and H2BE76 are destabilized. **(Q)** PottsMPNN predictions for how pair energies change upon the *H2B-D51H* mutation. Structural analysis of the neighboring residues highlights that interactions with H2AR71 and H2BH49 are destabilized. The structural visualizations were made using PDB: 2CV5.

To benchmark the sensitivity of the intracellular staining assay, we first examined canonical histone mutations with well-characterized biochemical effects. Expression of *H3*.*1-K27M* resulted in a strong reduction of global H3K27me3, while *H3*.*3-K36M* expression reduced global H3K36me3 levels. Both effects were detected by flow cytometry and confirmed by immunoblot (**Supplementary Figure 7A-B**), establishing that the screening platform reliably captures global chromatin perturbations associated with histone mutations. Validation experiments were performed by expressing individual histone mutations in MSCs followed by intracellular staining, flow cytometry, and quantification of PTM levels normalized to total histone H3 (see Methods and **Supplementary Figure 7C**).

#### Enhancer- and promoter-associated chromatin marks

Screening for H3K4me1 identified several mutations that strongly altered enhancer-associated chromatin. *H4-S1F* was associated with a pronounced increase in H3K4me1 signal, whereas *H4-K8N* and *H4-R17H* significantly reduced this mark (**Figure 3B-C**). Phosphorylation of H4S1 has been implicated in evolutionarily conserved mitotic regulation,^33^ and its disruption may alter chromatin accessibility at enhancer elements. H4K8 is normally acetylated in a manner that correlates with H3K27ac at active enhancers;^34^ substitution with asparagine eliminates this acetylation capacity, potentially contributing to reduced H3K4me1. H4R17 participates in a salt-bridge interaction with the MLL3 complex,^35^ and its mutation to histidine likely impairs enhancer methyltransferase recruitment.

Analysis of promoter-associated H3K4me3 revealed complementary perturbations. *H2A-R29Q* and *H3*.*3-R2G* significantly increased global H3K4me3 levels, while the well-characterized *H3*.*3-K4M* produced strong depletion of this modification (**Figure 3D-E**). Methylation of H2AR29 by PRMT6 has been linked to transcriptional repression,^36^ suggesting that loss of this modification site may derepress promoter activity. The relationship between H3R2 methylation and H3K4me3 is complex, with both cooperative and antagonistic interactions reported depending on cellular context and methylation state.^37–40^ *H3*.*3-R2G* emerged as a top hit that increased global H3K4me3, a finding confirmed by individual validation (**Figure 3D-E**). This result is consistent with a model in which loss of the H3R2 methylation site relieves inhibition of H3K4 methyltransferase activity, though alternative regulatory mechanisms described in other contexts cannot be excluded. As expected, *H3*.*3-K4M* markedly reduced global H3K4me3, in agreement with prior observations in preadipocyte systems^27^ (**Figure 3D-E**).

#### Heterochromatin-associated marks

We next examined constitutive and facultative heterochromatin. Several mutations, including *H2A-E56K* and substitutions at the conserved H3Q5 residue (*Q5H* and *Q5P*), increased global H3K9me3, suggesting a shift toward constitutive heterochromatin (**Figure 3F-G**). H3Q5 is a known site of serotonylation that cooperates with H3K4me3 to promote transcriptional activation through WDR5 recognition.^41–43^ These findings suggest that elimination of this N-terminal modification site favors heterochromatin accumulation. *H3*.*3-Q5P* also reduced global H3K4me3 (**Figure 3D**), consistent with a coordinated shift from active to repressive chromatin states. Analysis of facultative heterochromatin marked by H3K27me3 identified *H2A-S122N* and *H3*.*3-Q5H* as mutations that enhanced this modification, whereas *H3-K27M* produced the expected strong depletion of H3K27me3 (**Figure 3H-I**). H2AS122 is phosphorylated in response to DNA damage and has been implicated in chromosome segregation in yeast,^44,45^ suggesting a potential link between genome-maintenance pathways and PRC2-mediated repression.

#### Transcription-associated chromatin

Screening for H3K36me3 confirmed that both *H3*.*1-K36M* and *H3*.*3-K36M* strongly reduced global levels of this transcription-associated mark (**Figure 3J-K**), consistent with previous studies.^9,13^ In contrast, *H2A-R29Q, H2A-K75N*, and *H3*.*3-N108S* significantly increased H3K36me3. Acetylation of H2AK75 has been predicted to enhance nucleosomal DNA accessibility,^46^ and its loss through mutation may favor a shift toward transcription-associated chromatin states via increased H3K36me3. *H2A-P117T*, located near the PRC1-associated ubiquitination site H2AK119,^47^ also increased H3K36me3. Among H3 mutations, *H3*.*3-R63C* reduced global H3K36me3. H3R63 is a conserved sprocket arginine that directly contacts nucleosomal DNA and can be mono- or dimethylated.^48^ In yeast, substitution of this residue impairs binding of the FACT complex subunit Spt16, indicating a role in transcription-coupled nucleosome dynamics.^49,50^ Conversely, *H3*.*3-N108S* resulted in elevated global H3K36me3; this mutation has been shown to trap Spt16 at the 3’-end of highly expressed genes, disrupting normal transcriptional elongation dynamics.^51,52^ These opposing effects demonstrate that distinct classes of oncohistones can perturb transcription-associated chromatin through multiple mechanistically distinct pathways.

#### Global chromatin accessibility

Because histone modifications can influence nucleosome stability and higher-order chromatin organization, we next asked whether oncohistones also perturb global chromatin accessibility. We adapted a pooled ATAC-see^53,54^ strategy in which chromatin accessibility is measured by fluorescent Tn5 transposase labeling, followed by flow cytometry and barcode sequencing (**Figure 3L, Supplementary Figure 7D**). To control for cell-cycle-dependent variation, cells were gated in G1 phase; those within the upper or lower ∼5% of ATAC-see signal were isolated by FACS sorting at passages 1, 4, and 7, and only mutations that overlapped across all three passages were designated as high-confidence hits. Assay sensitivity and dynamic range were validated by treatment with the histone deacetylase inhibitor panobinostat,^55^ which produced a robust increase in histone acetyl-lysine and chromatin accessibility signal (**Supplementary Figure 7E-F**).

The ATAC-see screen identified mutations across all four core histone families that reproducibly altered global accessibility. *H2B-E76K, H4-R92T*, and *H3*.*3-K36M* significantly increased accessibility, whereas *H3*.*3-K4M* and *H3*.*3-G34L* reduced it (**Figure 3M-N**). *H3*.*3-K36M* emerged as the strongest accessibility-promoting hit and was validated individually (**Figure 3N**). Together with *H2B-E76K and H4R92T*, it represents a class of oncohistone mutations jointly associated with increased proliferation, impaired differentiation, and elevated global chromatin accessibility. *H3*.*3-E97K*, recently reported to inhibit PRC2 activity and histone H1 loading *in vitro*,^56^ also significantly increased accessibility, providing the first cellular-level evidence supporting those biochemical findings. Conversely, *H3*.*3-G34L* reduced accessibility, potentially reflecting increased H3K27me3 deposition as previously observed in HeLa cells.^57^

#### Computational modeling of nucleosome stability

Several accessibility-promoting mutations mapped to residues at histone-histone interfaces within the nucleosome core, suggesting that perturbation of nucleosome architecture may underlie these effects. To investigate the relationship between chromatin accessibility and intrinsic nucleosome stability, we used PottsMPNN, a graph-based neural network that models the sequence-energy landscape of protein structures,^58^ to computationally predict the effects of oncohistone mutations on mononucleosome stability energy (**Figure 3O**). This analysis focused on residues with observable electron density in the nucleosome crystal structure; most of the residues within disordered histone tails were excluded, as they had minimal predicted effects on stability. PottsMPNN computes pairwise interaction energies for neighboring residues, enabling identification of specific nucleosome structural contacts disrupted by a given mutation. To validate how well PottsMPNN models the energy landscape of the nucleosome, we compared its predictions to experimental nucleosome stability measurements^12^ and observed a moderately strong correlation (Pearson *r* = 0.51; **Supplementary Figure 7G**).

We focused on two accessibility-promoting hits: *H4-R92T* and *H2B-D51H* (**Figure 3M-N**). PottsMPNN predicted that *H4-R92T* disrupts interactions with H4D68 and H2BE76, both of which likely form favorable contacts with H4R92 in the nucleosome structure (**Figure 3P**). For *H2B-D51H*, the model predicted disruption of interactions with H2AR71 and H2BH49, again consistent with nucleosome structural context (**Figure 3Q**). However, PottsMPNN stability predictions did not explain all accessibility changes observed in the screen, suggesting that some mutations alter accessibility through mechanisms beyond mononucleosome destabilization, such as altered chromatin adaptor, reader, or remodeler recruitment.

Together, these results demonstrate that cancer-associated histone mutations can reprogram global chromatin accessibility in a residue- and structure-dependent manner. Mutations that enhanced accessibility frequently localized to interfaces critical for nucleosome stability, particularly the H2B-H4 interface. Computational modeling further revealed that accessibility-promoting mutations destabilize the nucleosome particle. Although calculated at the single-nucleosome level, such structural perturbations are expected to be amplified when mutant histones are deposited genome-wide, linking intrinsic nucleosome destabilization to the global chromatin remodeling observed in oncohistone-expressing cells.

### H3.3-Q5H is a context-dependent oncohistone that accelerates lung cancer growth via remodeling of active and repressive chromatin

Among the mutations identified in the pooled functional and chromatin screens, H3Q5 substitutions emerged as a distinctive class that concomitantly reduced active chromatin marks and increased repressive modifications in mouse mesenchymal stem cells (**Figure 3D-I**). We investigated whether Q5 mutations exhibit tumor-type specificity and promote growth in their cancer context of origin.

Because Q5 is conserved between canonical H3.1/2 and the replication-independent variant H3.3, we compiled mutation frequencies (n = 33) across all H3-encoding genes and identified three recurrent substitutions at this position: Q→H, Q→P, and Q→L (**Figure 4A**). These recurrent sub-stitutions underscore the functional importance of the conserved glutamine residue. Q5 mutations were most frequently observed in non-small cell lung cancer (NSCLC), which accounted for 24% of tumors harboring a Q5 substitution, followed by colorectal (12%) and ovarian (9%) cancers (**Figure 4A**). Tumors carrying Q5 mutations exhibited significantly lower median tumor mutational burden (TMB) than tumors with other histone mutations, consistent with positive selection and a potential driver role rather than a passenger event in a hypermutated background (**Figure 4B**).

**Figure 4:**
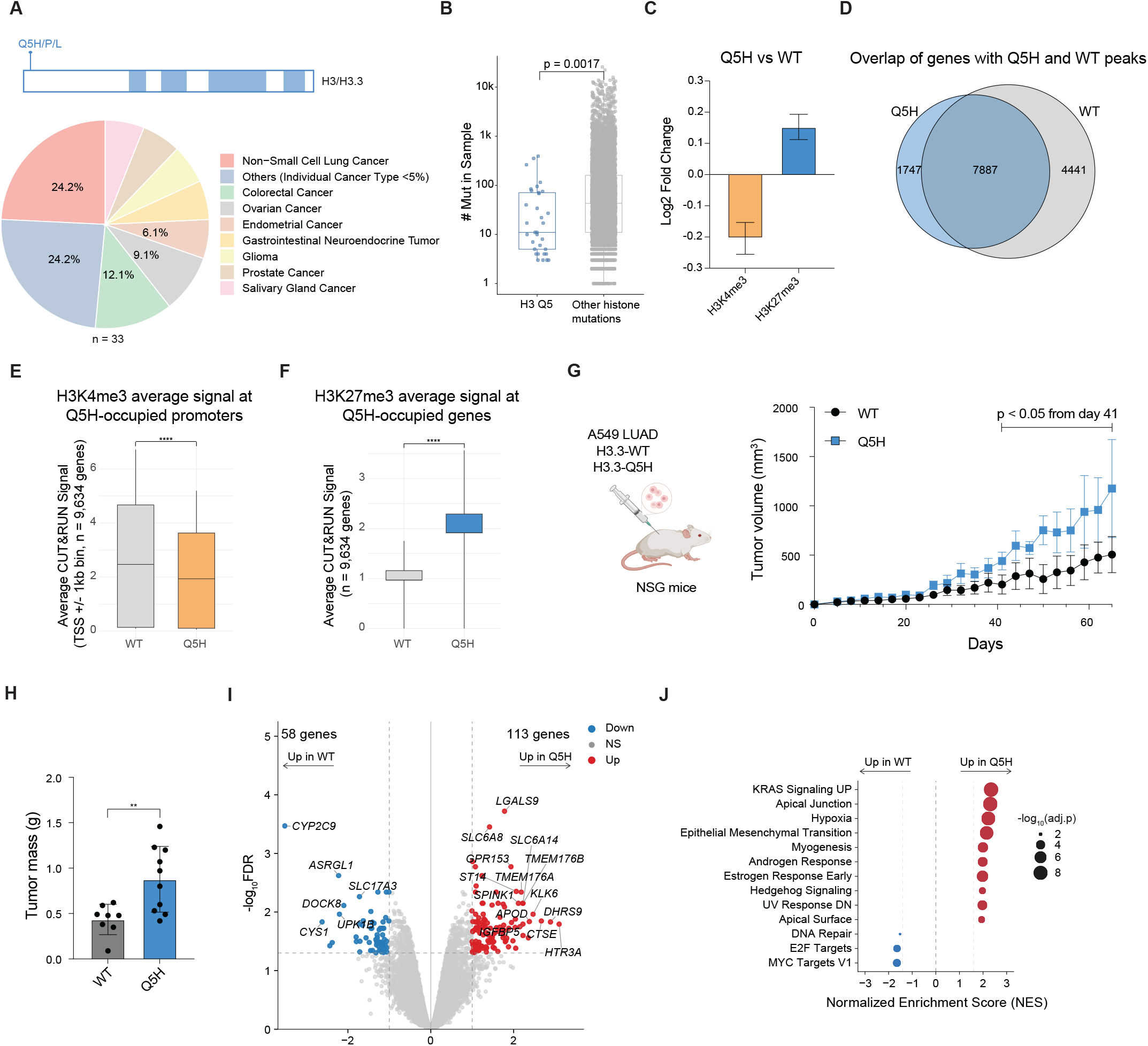
*H3*.*3-Q5H* oncohistone accelerates lung cancer growth and reshapes repressive chromatin landscapes. **(A)** Cancer type distribution for H3/H3.3 Q5 hotspot mutations (n = 33). Shaded area indicates histone fold domains. **(B)** Comparison of mutation numbers in tumor samples harboring Q5 mutations versus other histone mutations. **(C)** Plots of log_2_(fold-change) of H3.3-Q5H global histone PTM ICS data in A549 (normalized to EV; all values shown *P*-value < 0.05). **(D)** Venn diagram of genes with *H3*.*3-Q5H* and *H3*.*3-WT* CUT&RUN peaks. **(E)** Quantifications of H3K4me3 average CUT&RUN signal at H3.3-Q5H-occupied promoters (9,634 genes), TSS +/-1kb binned, non-parametric two-tailed Wilcoxon test, (****) indicates *P*-value < 0.0001. **(F)** Quantifications of H3K27me3 average CUT&RUN signal at H3.3-Q5H occupied promoters (9,634 genes). **(G)** Schematic of subcutaneous modeling of A549 cells expressing *H3*.*3-WT* or *H3*.*3-Q5H*. Tumor volume growth (mm^3^) monitored using caliper measurement across 65 days. **(H)** (Harvested tumor mass in grams (g) of subcutaneous tumors expressing *H3*.*3-WT* or *H3*.*3-Q5H*. **(I)** Volcano plot of differentially expressed genes measured by RNA-Seq in A549 subcutaneous tumors (Q5H vs WT), n = 4 replicates. Genes significantly upregulated in *Q5H* are shown in red (log_2_(fold-change) > 1 and -log_10_FDR > 1.3, n = 113 genes). Genes significantly upregulated in *WT* are shown in blue (log_2_(fold-change) < -1 and -log_10_FDR > 1.3, n = 58 genes). **(J)** Gene set enrichment analysis (GSEA) displaying the most significantly enriched MSigDB Hallmark gene sets in human A549 subcutaneous tumors expressing *H3*.*3-WT* or *H3*.*3-Q5H*.

Given that Q→H was the most frequent Q5 substitution in our pan-cancer analysis (**Figure 1E**), we expressed HA-tagged H3.3-Q5H in A549, a human lung adenocarcinoma cell line harboring an oncogenic *KRAS* mutation (*G12S*) (**Supplementary Figure 8A**). Intracellular staining analysis confirmed that *H3*.*3-Q5H* expression in this cell line is associated with reduced global H3K4me3 and increased global H3K27me3 (**Figure 4C**), consistent with the chromatin signature observed by intracellular staining in mouse mesenchymal stem cells (**Figure 3E, 3I**). CUT&RUN profiling of early-passage cells further confirmed genome-wide alterations of these marks, with widespread loss of promoter-associated H3K4me3 and expansion of repressive H3K27me3 domains (**Supplementary Figure 8B-E**).

To determine the genomic deposition profile of H3.3-Q5H, we compared the set of genes with HA CUT&RUN peaks between Q5H-and WT-expressing cells and found extensive overlap (**Figure 4D**). Peaks displayed similar distributions across genomic features, indicating that the N-terminal tail substitution does not substantially alter interactions with histone chaperones or the global genomic deposition profile (**Supplementary Figure 8F)**. Nevertheless, at H3.3-Q5H-occupied genes, H3K4me3 was markedly reduced at promoter regions while H3K27me3 was increased across gene bodies (**Figure 4E-F**), suggesting that Q5H exerts its chromatin effects locally at sites of incorporation.

We next asked whether this epigenomic reprogramming alters cellular fitness. In contrast to the proliferation defects observed in mouse mesenchymal stem cells (**Figure 2C**), A549 cells expressing *H3*.*3-Q5H* displayed a significant proliferative advantage over WT-expressing controls in an *in vitro* growth competition assay (**Supplementary Figure 8G**), indicating that the functional consequence of *Q5H* is strongly context-dependent. To determine whether this advantage extends to tumor growth *in vivo*, we injected A549 cells expressing *H3*.*3-WT* or *H3*.*3-Q5H* subcutaneously into immunodeficient NSG mice (**Figure 4G**). Tumor growth kinetics diverged significantly beginning at day 41 post-injection, with *H3*.*3-Q5H* tumors exhibiting accelerated growth (**Figure 4G**). Tumor mass at endpoint was significantly greater in the Q5H cohort than in the WT cohort (**Figure 4H**), demonstrating that this mutation confers a growth advantage *in vivo*. Sustained transgene expression was confirmed by anti-HA immunohistochemistry on harvested tumors (**Supplementary Figure 8H**).

To define the transcriptional programs underlying Q5H-driven tumor growth, we performed RNA sequencing on harvested tumors (**Figure 4I**). Differential expression analysis identified upregulation of multiple genes implicated in lung cancer progression. Among the most strongly induced transcripts was *HTR3A*, which has been linked to proliferation through FOXH1, Wnt3A signaling, and ERK activation.^59,60^ *TMEM176A* and *TMEM176B* were also significantly upregulated, suggesting coordinated activation of pro-tumorigenic programs including KRAS signaling.^61,62^ Additional upregulated genes included members of the SLC6 transporter family (*SLC6A8, SLC6A14*) and *KLK6*, a kallikrein previously implicated in lung adenocarcinoma progression and immune evasion.^63^ Gene set enrichment analysis further showed enrichment of KRAS signaling, epithelial-mesenchymal transition (EMT), hypoxia, and apical junction programs in Q5H tumors, whereas MYC target, E2F target, and DNA repair programs were enriched in WT tumors (**Figure 4J**). Intriguingly, this inverse relationship may suggest a form of oncogenic buffering, where simultaneous hyperactivation of KRAS and MYC signaling reaches antagonistic levels that compromise cell viability, requiring tumors to maintain a balance between these programs.

Collectively, these findings establish H3.3-Q5H as a novel context-dependent oncohistone that promotes lung adenocarcinoma growth through coordinated suppression of promoter-associated H3K4me3 and expansion of repressive H3K27me3 domains. These chromatin changes drive transcriptional reprogramming associated with KRAS signaling hyperactivation (beyond the *KRAS*^G12S^ mutation) and EMT programs. The low TMB of Q5-mutant tumors in NSCLC patients further supports a driver rather than passenger role for this cancer mutation.

## Discussion

Our systematic functional interrogation of 303 cancer-associated histone mutations establishes several principles governing how oncohistones reshape chromatin biology and cellular behavior, and provides a resource framework for continued mechanistic and therapeutic exploration.

A central finding of our pancancer analysis is that histone mutations are not randomly distributed across the genome. Despite encoding nearly identical or in many cases identical proteins, paralogous histone genes harbor distinct mutational hotspots and display divergent cancer-type enrichment patterns. This observation cannot be explained by protein sequence alone and instead implicates gene-level features such as expression timing, replication-coupled versus replication-independent deposition, chromatin environment at the gene locus, or locus-specific regulatory elements as determinants of mutational selection in cancer. These findings argue that histone genes should not be treated as functionally interchangeable in cancer genomics, even when their encoded proteins are nearly identical. Furthermore, the strong biases in amino acid substitution class, particularly the enrichment of arginine loss and charge-reversing substitutions in low-TMB contexts, support the view that many histone mutations arise under positive selection rather than as byproducts of hypermutation. Together, these patterns expand the oncohistone concept well beyond the *H3K27M* and *H3K36M* paradigm and suggest that a substantial fraction of the recurrent histone mutations identified in adult cancers may function as drivers or modifiers of tumor fitness.

Pooled screening across proliferation, differentiation, five histone PTMs, and chromatin accessibility revealed that functionally consequential mutations are not uniformly distributed across the nucleosome but instead cluster at specific structural features. Most strikingly, mutations at the H2B-H4 interface (*H2B-E76K, H4-R92T*) consistently promoted proliferation, blocked differentiation, and increased chromatin accessibility. These residues reside on different histone subunits yet form direct contacts within the nucleosome, and their convergence on shared phenotypes suggests that precise destabilization of histone-histone interactions can create a permissive chromatin state that broadly enhances cancer cell fitness. This convergence also argues against simple overexpression toxicity. If bulk histone over-production were responsible, structurally unrelated mutations would be expected to produce similar effects. Instead, phenotypic consequences track with disruption of defined nucleosome contacts, supporting a structure-based mechanism. Further supporting this model, computational analysis with PottsMPNN predicted how some accessibility-promoting mutations destabilize core interfaces. However, stability predictions did not account for all observed chromatin accessibility changes, indicating that additional mechanisms also contribute to the global chromatin effects of oncohistones.

Our findings extend prior work demonstrating that *H2B-E76K* promotes growth in mammary epithelial cells^22^ by showing that this phenotype generalizes to mesenchymal stem cells, is mirrored by its structural partner *H4-R92T*, and is accompanied by concomitant effects on differentiation and chromatin accessibility. They also complement the structural predictions of Nacev et al.^14^ by providing direct functional evidence that nucleosome-destabilizing mutations at predicted hotspots produce measurable chromatin and cellular phenotypes. Systematic histone mutagenesis studies in yeast similarly established that residues at histone-histone interfaces and DNA contact points are critical for fitness and transcriptional regulation,^64,65^ and our mammalian data confirm this principle across eukaryotes. However, we also observe phenotypes such as the strong proliferative advantage conferred by *H4-R92T* and the differentiation-blocking activity of *H2A-K74N* that have no clear counterpart in yeast screens, likely reflecting the additional regulatory complexity of mammalian chromatin.

Among the unexpected results of our screens, H4 mutations were broadly pro-proliferative, a finding not anticipated from prior literature, which has focused predominantly on H3 variants. Conversely, H3.3 was generally intolerant of perturbation outside established hotspots such as K27 and K36, suggesting that the H3.3 variant is under tighter functional constraint in mesenchymal stem cells than its canonical H3 counterpart. These observations highlight that systematic, unbiased screening can reveal biology obscured by candidate-based approaches focused on known oncogenic residues.

Our chromatin-based screens identified 51 mutations across all histone families that globally alter histone PTM landscapes or chromatin accessibility, substantially expanding the oncohistone functional repertoire to include underexplored mutations in H2A, H2B, and H4. These effects span enhancer-associated (H3K4me1), promoter-associated (H3K4me3), heterochromatin-associated (H3K9me3, H3K27me3), and transcription-associated (H3K36me3) chromatin states. Several findings highlight mechanistic diversity within this landscape. *H3*.*3-R63C* and *H3*.*3-N108S* both perturb the FACT complex yet produce opposing effects on H3K36me3, demonstrating that oncohistones can converge on shared pathways while generating divergent chromatin outcomes. H3Q5 substitutions coordinately reduce H3K4me3 and increase H3K9me3 and H3K27me3, revealing a mutation class that simultaneously suppresses active chromatin and expands repressive domains. This dual activity was not previously described outside the canonical K→M paradigm. The high-throughput assays developed here, pooled intracellular staining of PTMs and pooled ATAC-see, provide a scalable, single-cell-resolution strategy for mapping chromatin perturbations across large mutational libraries and should be broadly applicable to other classes of chromatin-associated cancer mutations.

A clinically-relevant finding is the identification and validation of *H3*.*3-Q5H* as a context-dependent oncohistone that accelerates lung adenocarcinoma growth *in vivo*. Expression of *H3*.*3-Q5H* in mesenchymal stem cells suppressed cellular proliferation, impaired differentiation, and shifted the chromatin landscape toward repressive states. In contrast, expression of *H3*.*3-Q5H* in A549 lung adenocarcinoma cells conferred a significant proliferative advantage and accelerated tumor growth *in vivo*. CUT&RUN profiling confirmed suppression of promoter-associated H3K4me3 and expansion of repressive H3K27me3 domains at sites of mutant histone deposition, while RNA sequencing revealed activation of KRAS signaling and EMT programs. The enrichment of KRAS pathway transcripts is notable given that A549 cells harbor an endogenous oncogenic *KRAS*^*G12S*^ allele, raising the possibility that Q5H amplifies a pre-existing oncogenic program rather than establishing one *de novo*. Mechanistically, Q5 is the site of serotonylation (H3Q5ser), which cooperates with H3K4me3 to recruit WDR5 and promote transcription.^41,42^ Substitution to histidine eliminates this modification site, providing a plausible basis for the observed H3K4me3 reduction and shift toward repressive chromatin. More broadly, the finding that *Q5H* promotes growth specifically in lung adenocarcinoma, the cancer type in which it is most frequently observed, parallels the tissue specificity of *H3K27M* in glioma and *H3K36M* in chondroblastoma,^9,10^ and further underscores the importance of assessing oncohistone function in disease-relevant models.

Because oncohistones globally alter histone PTM landscapes and chromatin accessibility, they have the potential to modulate sensitivity to chromatin-targeted therapies, including inhibitors of histone methyltransferases, demethylases, adaptor, or reader proteins.^66,67^ The CHANCLA-303 library and pooled screening platforms established here are directly compatible with drug-sensitivity mapping, enabling systematic identification of synthetic dependencies created by specific chromatin perturbations. Furthermore, global chromatin remodeling can influence antigen presentation, transposable element de-repression, and immune checkpoint regulation.^68,69^ Extension of oncohistone screening to immune coculture systems or immunocompetent *in vivo* models may reveal how specific histone mutations shape tumor-immune interactions and modulate response to immunotherapy.

This study provides the most comprehensive functional characterization of cancer-associated histone mutations to date, integrating pan-cancer genomic analysis, pooled phenotypic and chromatin-based screening, computational modeling, and *in vivo* validation into a unified resource. Our findings demonstrate that oncohistones follow evolutionary and structural constraints, converge on specific nucleosome interfaces to reprogram chromatin and cancer cell fate, and can function as context-dependent drivers of tumor growth. The experimental platforms and functional atlas established here provide a foundation for dissecting the mechanistic and therapeutic implications of the expanding oncohistone landscape.

## LIMITATIONS OF THE STUDY

This study provides a systematic functional atlas of cancer-associated histone mutations in mammalian cells, but several limitations should be considered.

First, all pooled screens were performed using lentiviral cDNA overexpression rather than endogenous gene editing. Although this approach enables scalable comparison across hundreds of mutations under controlled expression conditions, it does not fully recapitulate native allelic dosage or cell cycle-coupled chromatin deposition dynamics. In theory, studies using endogenous editing of histone loci could allow for better assessment of native allele-specific and dosage-dependent effects. However, physiologically accurate editing of endogenous histone loci presents major technical challenges in part due to the multi-copy clustered nature of histone gene families and their virtually identical sequence composition, which could exacerbate off-target effects. It is worth noting that canonical oncohistones, including H3K27M and H3K36M, exhibit a genetically dominant oncogenic phenotype, which can be modeled appropriately using cDNA-based expression systems.

Second, our recurrence filter (≥3 patients) was necessary to enrich for likely functional mutations but may exclude rare yet potent drivers present in only one or two patients.

Third, functional screening experiments were conducted primarily in mesenchymal stem cells, chosen for their differentiation plasticity and established utility in classic oncohistone studies. While this system enabled identification of proliferation, differentiation, and chromatin phenotypes in a controlled context, our own data demonstrate that histone mutation consequences are strongly lineage-dependent, as illustrated by the divergent phenotypes of *H3*.*3-Q5H* in MSCs versus lung adenocarcinoma cells. Extending systematic screening to additional tissue-specific and genetically defined models will be necessary to capture the full spectrum of context-dependent oncohistone behavior.

Fourth, the chromatin profiling approaches employed here quantify global shifts in histone modifications and accessibility but do not resolve locus-specific chromatin architecture at single-cell or three-dimensional levels. CUT&RUN analyses for selected mutations provide genome-wide mapping in defined contexts, but higher-resolution approaches, including single-cell chromatin profiling and chromosome conformation assays, will be needed to clarify how nucleosome-scale perturbations propagate to domain-level and compartment-level reorganization.

Finally, *in vivo* validation was performed using subcutaneous transplantation into immunodeficient mice. While these experiments demonstrate that recurrent histone mutations can enhance tumor growth in their cancer context of enrichment, they do not address effects on immune surveillance, metastatic dissemination, or therapeutic response. Given the global chromatin remodeling induced by oncohistones, future studies incorporating immunocompetent models, orthotopic systems, and therapeutic perturbations will be essential to define their roles in tumor-microenvironment interactions and treatment sensitivity.

## Methods

### Plasmids and Cloning

Human histone sequences were acquired from UniProt, and cDNA gene fragments were synthesized by Twist Bioscience. The backbone plasmid was modified from pCDH-EF1α-MCS-IRES-Puro (System Biosciences, CD532A-2). The pooled cDNA library was digested with EcoRI and BamHI restriction enzymes (New England BioLabs), then ligated to the digested backbone using T4 ligase (New England BioLabs). The purified ligation product was electroporated into Lucigen Endura ElectroCompetent Cells (Biosearch Technologies) before being plated on LB-carbenicillin plates (Teknova, L5010). Library representation was assessed via serial dilution plating. Ten random colonies from these serial dilution plates were picked to assess the fidelity of library cloning. Bacteria were scraped from the plates and incubated in LB-carbenicillin for 2 h at 37 °C. DNA was purified using QIAGEN Maxiprep Kit, following the manufacturer’s protocol. A complete list of cDNA and primer sequences used in this study is provided in **Supplementary Table 2**.

### Cell Culture

Human HEK293T (ATCC) cells were maintained in DMEM (Corning) supplemented with 10% FBS (Gibco), penicillin (100 U/mL, Gibco), streptomycin (100 µg/mL, Gibco), and plasmocin (5 µg/mL, InvivoGen). A549 (ATCC) cells were maintained in DMEM (Corning) supplemented with 10% FBS (Gibco), penicillin (100 U/mL, Gibco), and streptomycin (100 µg/mL, Gibco). Mouse C3H/10T1/2 mesenchymal stem cells (ATCC) were maintained in DMEM (Corning) supplemented with 10% FBS (Gibco), 1% GlutaMAX (Gibco), streptomycin (100 µg/mL, Gibco), and penicillin (100 U/mL, Gibco). Cell lines were confirmed to be free of mycoplasma contamination and cultured at 37°C and 5% CO_2_.

### Lentiviral Transduction

Lentivirus was produced in HEK293T cells (ATCC) transfected with psPAX2 (Addgene, 12260) and VSV-G (Addgene, 14888) lentiviral plasmids using Lipofectamine 2000 (Thermo Fisher Scientific). Virus was collected 48 and 72 h after transfection and filtered through a 0.45 µm filter. Viral supernatants were applied directly to target cells with the addition of 1 mg/mL polybrene (Millipore-Sigma). Transduced cells were then selected with puromycin (Thermo Fisher Scientific).

### Pooled cDNA Screen Conditions and Analysis

To ensure that most cells had a single lentivirus integration event, we determined the volume of viral supernatant that would achieve an MOI of ∼0.3 upon infection of a population of mouse mesenchymal stem cells. Briefly, cells were plated at a concentration of 2.5 × 10^5^ cells per well in 12-well plates along with increasing volumes of library pool viral supernatant and polybrene (1 mg/mL, EMD Millipore). Viral infection efficiency was determined by the percentage of RFP+ cells assessed by flow cytometry on an LSRFortessa (BD Biosciences) instrument 72 h post-infection. A total of 7.5 × 10^5^ cells were infected for each replicate of the screen. All screens (proliferation, differentiation, histone PTM ICS, and ATAC-see) were infected using the same procedure.

In the proliferation z-score analysis, for each passage, we computed the average read count and coefficient of variation (CV) for every unique lentiviral barcode across three replicates. Barcodes were filtered to retain only those with stable abundance profiles, defined as a median CV < 0.65 across the twelve passages. For each retained barcode, we constructed a passage-wise abundance matrix and normalized the values using row-wise z-score transformation. We then fitted a simple linear regression model to get the linear slope, where z_i_ represents the z-score at passage p_i_∈[0,1,…,11], β_1_ is the slope (rate of change in enrichment), and ϵ_i_ is the residual error.

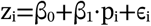

In the intracellular staining (ICS), differentiation, and ATAC-see screens, for each passage, we computed the average read count and coefficient of variation (CV) for every unique lentiviral barcode across three replicates. Barcodes were filtered to retain only those with stable abundance profiles, defined as a median CV < 0.65 across the passages. Differential abundance between high and low populations was quantified as the log_2_(fold-change) of mean percentage abundance. To identify reproducibly enriched lentiviral barcodes associated with the high or low population, hits were defined using percentile-based thresholds within each passage. The top 10% of lentiviruses with the highest log_2_(fold-change) values were classified as enriched hits, whereas the bottom 10% with the lowest log_2_(fold-change) values were classified as depleted hits. For depleted hits, absolute log_2_(fold-change) values were used for ranking and display purposes. Overlap between enriched or depleted hit sets across two passages was assessed by set intersection and visualized using two-set Venn diagrams.

### Next-Generation Sequencing

Genomic DNA was extracted using the DNeasy Blood & Tissue Kit (QIAGEN). We assumed that each cell contains approximately 6.6 pg of genomic DNA (gDNA). Therefore, deconvolution of the screen at 1000X coverage required sampling at least 300,000 cells x 6.6 pg of gDNA per cell. Library preparation was done by GENEWIZ NGS service using Platinum SuperFi II Master Mix and Equinox Master Mix. The library was sequenced by GENEWIZ NGS service using the NovaSeq X Plus platform. Raw reads were trimmed with Trimmomatic v0.36 and paired read 1 and read 2 were subsequently merged using the BBMerge tool from the BBMap software suite. A hash table count was performed on the merged reads to identify any region nested by the forward and reverse primer pair.

### Expression and Purification of Recombinant Tn5

Recombinant hyperactive Tn5 transposase was expressed and purified from bacteria (*E. coli*) following a modified protocol ^70,71^. Briefly, cells were grown to an optical density of ∼0.6 at 600 nm prior to induction. After induction and expression, cells were harvested by centrifugation and stored at –80 °C until use. Cell pellets were resuspended in freshly prepared lysis buffer (20 mM HEPES pH 7.6, 800 mM NaCl, 1 mM EDTA, 5 mM β-mercaptoethanol, 0.1% Triton X-100, 10% glycerol, and 1X protease inhibitor cocktail). Cells were lysed by sonication and clarified by centrifugation at 14,000 rpm for 15 min at 4 °C. The supernatant was incubated with 4 mL pre-equilibrated Ni-NTA agarose resin (Qiagen) for 4 hours at 4 °C with gentle rotation. Prior to use, the resin was washed and equilibrated with equilibration buffer (20 mM HEPES pH 7.6, 500 mM NaCl). Bound proteins were washed sequentially with wash buffer 1 (20 mM HEPES pH 7.6, 800 mM NaCl, 20 mM imidazole, 1 mM EDTA, 10% glycerol, 2 mM β-mercaptoethanol, 0.05% Triton X-100) and wash buffer 2 (20 mM HEPES pH 7.6, 150 mM NaCl, 30 mM imidazole, 1 mM EDTA, 5% glycerol). Proteins were eluted using elution buffer (20 mM HEPES pH 7.6, 150 mM NaCl, 1 mM EDTA, 5% glycerol, 300 mM imidazole, 3 mM β-mercaptoethanol). Purified Tn5 fractions were pooled and ion-exchanged using a HiTrap SP HP column connected to an ÄKTA pure chromatography system (Cytiva). Eluted fractions were analyzed by SDS-PAGE followed by Coomassie staining.

### Histone Extraction

Cell pellets were collected by centrifugation and washed once with ice-cold 1X PBS (Gibco) supplemented with protease inhibitors (cOmplete EDTA-free, Roche). Pellets were either processed immediately or stored at –80 °C. Pellets were resuspended in hypotonic buffer (10 mM Tris-HCl pH 8.0, 1 mM KCl, 1.5 mM MgCl_2_, 1 mM PMSF, 1 mM DTT, and protease inhibitors) and incubated on ice for 15 min. NP-40 (Sigma-Aldrich) was then added to a final concentration of 0.3%, followed by an additional 5-min incubation on ice. Nuclei were pelleted by centrifugation at 10,000 rpm for 5 min at 4 °C. The nuclear pellet was resuspended in 200 µL of 0.4 N sulfuric acid (H_2_SO_4_; Sigma-Aldrich) and incubated with rotation at 4 °C for 2 h to overnight. Acid-insoluble material was removed by centrifugation at 13,000 rpm for 10 min at 4 °C. Proteins in the supernatant were precipitated by the addition of 66 µL of 100% trichloroacetic acid (TCA; Sigma-Aldrich), followed by gentle inversion and incubation on ice for 30 min. Precipitated histones were collected by centrifugation at 13,000 rpm for 10 min at 4 °C, washed with 1 mL of ice-cold acetone, and air-dried. Dried protein pellets were resuspended in ultrapure water and subjected to brief sonication in the Bioruptor Pico system (Diagenode). Histone concentration was quantified using a Qubit Flex Fluorometer (Thermo Fisher Scientific).

### Immunoblot Analysis

Proteins were resolved by SDS-PAGE and transferred to nitrocellulose membrane. Membranes were blocked in 5% non-fat dry milk in 1X TBS-T for 30 min at room temperature and probed with primary antibody overnight at 4 °C. After three washes with 1X TBS-T, membranes were incubated with secondary antibody for 1 h at room temperature. Antibodies used for immunoblotting: anti-HA tag (CST, C29F4), anti-H3K27me3 (CST, C36B11), anti-H3K36me3 (abcam, ab9050), rabbit anti-acetyl-lysine mAb (PTM Bio, PTM-105RM), mouse anti-rabbit IgG (light-chain specific) mAb (HRP conjugate) (CST, D4W3E). After three washes with 1X TBST, membranes were incubated in enhanced chemiluminescence (ECL) substrate (Thermo Fisher Scientific), and signals were captured using a ChemiDoc imaging system (Bio-Rad).

### RT-qPCR Analysis

RNA was extracted using TRIzol reagent (Invitrogen), following the manufacturer’s protocol. cDNA was synthesized from up to 2 µg of total RNA in 20 µL reactions using the High-Capacity cDNA Reverse Transcription Kit (Thermo Fisher Scientific) according to the manufacturer’s instructions. Quantitative PCR was performed using commercially available probes from TaqMan Gene Expression Assays: *Cebpa* Mm00514283_s1, *Pparg* Mm00440940_m1, *Actb* Mm02619580_g1 on a QuantStudio 6 Pro Real-Time PCR system (Applied Biosystems).

### Adipocyte Differentiation

Mouse C3H/10T1/2 mesenchymal stem cells were seeded in Dulbecco’s Modified Eagle Medium (DMEM; Thermo Fisher Scientific) supplemented with 10% fetal bovine serum (FBS; Gibco) and allowed to reach 100% confluency. Confluent monolayers were then exposed to an adipogenic induction medium comprising 0.5 mM isobutylmethylxanthine (IBMX; Sigma-Aldrich), 1 µM dexamethasone (Sigma-Aldrich), 5 µg/mL insulin (Sigma-Aldrich), and 5 µM troglitazone (Sigma-Aldrich), prepared in DMEM + 10% FBS. After 48 hours of induction, the medium was replaced with the maintenance medium containing only 5 µg/mL insulin in DMEM + 10% FBS. The maintenance medium was refreshed every 2-3 days thereafter.

### High-Throughput Sequencing and Computational Analysis

High-throughput sequencing libraries were prepared using Platinum SuperFi II Master Mix and Equinox Master Mix. Sequencing was performed on the NovaSeq X Plus platform (Illumina). Raw sequencing reads were quality-trimmed with Trimmomatic (v0.36) and paired-end reads (read 1 and read 2) were subsequently merged using BBmerge from the BBMap software suite. A hash table-based analysis was conducted on merged reads to identify regions flanked by the forward and reverse primer pairs.

### Computation of Nucleosome Residue Distance Matrix

Atomic coordinates were retrieved from the high-resolution nucleosome structure (PDB: 2CV5). Distances between all possible amino acid pairs were calculated using the Euclidean norm of Cα atom positions (equation shown below). In cases where residues were present in multiple chains, the minimum pairwise distance among all chain combinations was recorded. Residues with missing coordinates or lacking Cα atoms in the structure were excluded from the computation.

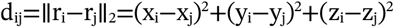

### Intracellular Staining (ICS)

Target cell lines were collected by trypsinization, fixed and permeabilized using the Foxp3/Transcription factor staining buffer set (CST 43481S) for 1 h at 4 °C. Cells were pelleted, washed with permeabilization buffer, then incubated with primary antibody for 1 h at room temperature. Cells were washed twice then incubated with fluorescent secondary antibody for 1 h at room temperature. Next, cells were washed twice in FACS buffer (10% FBS in PBS). Fluorescence was analyzed on a Celesta HST-1 (BD Biosciences) analytical flow cytometer. Data were analyzed using FlowJo (BD Life Sciences). The G1 cell population was gated using the DAPI signal. Normalized PTM staining MFI over total histone H3 staining MFI (FITC/APC) was used to calculate log_2_(fold-change) relative to empty-vector (EV) controls. Antibodies used for ICS were: anti-H3 (CST, #14269), anti-H3K4me1 (abcam, ab8895), anti-H3K4me3 (CST, C42D8), anti-H3K9me3 (abcam, ab8898), anti-H3K27me3 (CST, C36B11), anti-H3K36me3 (abcam, ab9050), anti-HA tag (CST, C29F4), mouse anti-rabbit IgG (light-chain specific) (HRP conjugate) (CST, D4W3E), goat anti-rabbit IgG (H+L)-Alexa Fluor Plus 488 (Thermo Fisher Scientific, A32731), DAPI (BD Pharmingen).

### BODIPY Staining

BODIPY™ FL C12 (D3822, Thermo Fisher Scientific) was diluted 1:2000 in Dulbecco’s Modified Eagle Medium (DMEM; Thermo Fisher Scientific) supplemented with 5 µg/mL insulin (Sigma-Aldrich), and incubated with differentiated C3H/10T1/2 cells at 37 °C for 30 min. After incubation, cells were washed twice with 1 mL pre-warmed FACS buffer (1X PBS, 5% FBS, 2 mM EDTA). Cells were then dissociated and gently resuspended. Samples were processed immediately by flow cytometry to quantify BODIPY fluorescence.

### ATAC-see

Tn5 transposase was recombinantly purified from *E. coli* cells following an established protocol ^71^. ATAC-see protocol was adapted and optimized from previously described protocols ^53,54^. Specifically, fluorescently labeled adaptor DNA was prepared by annealing equimolar ME-fw-ATTO488 and ME-rev[Phos] to generate a 45 µM duplex. Tn5 transposome was assembled by incubating equimolar purified Tn5 transposase with adaptor duplex for 1 hour at room temperature. Cells were washed and permeabilized with NP buffer (0.1% IGEPAL, 0.5% BSA, 1× TAPS), followed by a wash with cold 1X TAPS containing 0.5% BSA. For each reaction (∼500k cells), 500 µL of ATAC-see reaction mix was prepared by combining 330 µL nuclease-free water, 100 µL 5X TAPS buffer (50 mM TAPS pH 8.5, 25 mM MgCl_2_), 50 µL DMF, and 20 µL Tn5 transposome. Cells were incubated with 100 µL of the ATAC-see reaction mix at 37°C for 30 min. Reactions were stopped with two washes in 100 µL wash buffer (0.1% IGEPAL in 1X PBS). Cells were resuspended in FACS buffer containing DAPI (1 µg/mL final), incubated on ice for 15 min. Fluorescence signals from FITC channel and DAPI were quantified using flow cytometry. The G1 cell population was gated using the DAPI signal. Normalized ATAC-see probe MFI over DAPI MFI (FITC/DAPI) was used to calculate log_2_(fold change). For the sensitivity test, 500,000 C3H/10T1/2 cells were plated and treated with 10 nM panobinostat overnight, followed by ATAC-see protocol and flow cytometry analysis.

### Computational Prediction of Nucleosome Stability

PottsMPNN analysis was done using the code available on the GitHub repository, specifically the stability fine-tuned version of the model weights (https://github.com/KeatingLab/PottsMPNN). To predict specific interactions disrupted by mutations, we extracted the Potts model generated by the model using the nucleosome structure (PDB: 2CV5) and compared the pairwise energies made by the wildtype residue with those made by the mutant residue.^28^ We excluded interactions with a negligible (less than 0.25 a.u.) predicted energy difference from the plots. PyMOL v3.1 was used to visualize the structural context of selected mutant positions.

### RNA-Sequencing and Analysis

RNA was extracted from the dissected tumors following the manufacturer’s TRIzol Reagent protocol (Invitrogen, 15596026). Purified RNA was sequenced by Plasmidsaurus RNA-Seq service. Briefly, for sequencing library preparation, mRNA was converted into complementary DNA (cDNA) via reverse transcription and second-strand synthesis, followed by tagmentation, library indexing, and amplification. Illumina sequencing was used and 3’ end counting was used to capture differential gene expression. FastQ generation and demux was done with BCL Convert v4.3.6 and fqtk v0.3.1. Read-filtering used FastP v0.24.0: poly-X tail trimming, 3’ quality-based tail trimming, a minimum Phred quality score of 15, and a minimum length requirement of 50 bp. Alignment to the appropriate reference genome using STAR aligner v2.7.11 with non-canonical splice junction removal and output of unmapped reads. Coordinate sorting of BAM files using samtools v1.22.1. UMI based de-duplication: Removal of PCR and optical duplicates using UMICollapse v1.1.0. Mapping QC: Alignment quality metrics, strand specificity, and read distribution across genomic features using RSeQC v5.0.4 and Qualimap v2.3. Generation of comprehensive QC report using MultiQC v1.32. Gene-expression quantification using featureCounts (subread package v2.1.1) with strand-specific counting, multi-mapping read fractional assignment, exons and 3’-UTR as the feature identifiers, and grouped by gene_id. Final gene counts were annotated with gene biotype and other metadata extracted from the reference GTF file. Sample-sample correlations for sample-sample heatmap and PCA were calculated on normalized counts (TMM, trimmed mean of M-values) using Pearson correlation.

### CUT&RUN and Analysis

Nuclei were extracted from 500,000 cells using CUTANA Nuclei Extraction Buffer (Epicypher, 21-1026) according to the manufacturer’s instructions. CUT&RUN was subsequently performed using the CUTANA ChIC/CUT&RUN Kit (Epicypher, 14-1048). Extracted nuclei were conjugated to Concanavalin A beads and incubated with 2 µg of primary antibody overnight at 4°C. Primary antibodies used were: anti-H3K4me3 (EpiCypher, 13-0060), anti-H3K27me3 (EpiCypher, 13-0055). MNase conjugation and digestion were performed the following day. CUT&RUN fragments were purified using SPRI beads and quantified using 1X dsDNA High Sensitivity Reagents using Qubit (Thermo Fisher Scientific). Library preparation was performed using CUTANA CUT&RUN Library Prep Kit (Epicypher, 14-1001) using the manufacturer’s instructions. Libraries were quantified using Tapestation (Agilent). Sequencing was done using the Element AVITI24 to obtain >10 million 75bp paired-end reads at the Koch Institute BioMicroCenter. Raw sequencing reads were analyzed and converted to bigWig files using the nfcore/cutandrun pipeline (v3.2.2).^72^ Briefly, FASTQ files were quality checked using FastQC (v0.12.1). Reads were trimmed using TrimGalore (v0.6.6) and aligned to the human reference genome (hg38) and *E*.*coli* K12 spike-in genome using Bowtie2 (v2.4.4). Target reads were normalized according to spike-in levels. BAM files were sorted and indexed with SAMtools (v1.17), and duplicate reads were marked using Picard (v3.1.0). Any regions included in the CUT&RUN suspect list were blacklisted and excluded from downstream analyses.^73^ Peak calls were made relative to IgG control samples using MACS2 (2.2.7.1) in narrow peak mode (H3K4me3) or broad peak mode (H3K27me3, HA) filtering regions with q-value threshold of < 0.01. Annotation of genomic regions to genes was performed using the ChIPseeker R Bioconductor package (v1.46.1). Range-based heatmaps showing CUT&RUN signals over genomic regions were generated using Deeptools computeMatrix and plotHeatmap (v3.5.6) in either reference-point or scale-regions mode. CUT&RUN signal was binned using multiBigwigSummary (v3.5.6) using either the BED-file or bins mode. For the quantifications of average CUT&RUN signal, occupancy is defined by the presence of at least one H3.3-Q5H HA peak within the gene.

### *In vitro* Growth Competition Experiments

Cell lines transduced with oncohistone lentivirus (RFP+) were mixed with parental cell lines (RFP-) in 12-well plates at ∼30% RFP+/70% RFP-. Cells were monitored by flow cytometry over time using an LSRCelesta (BD Biosciences). Flow cytometry data were analyzed with FlowJo software (BD Biosciences). The percentage of RFP+ cells was normalized to their respective T0 time-point values.

### *In vivo* Tumorigenesis Assay

All animal experiments were performed in accordance with institutional guidelines and approved by the Committee on Animal Care (CAC) at the Massachusetts Institute of Technology. NSG mice (6-8 weeks, female sex-matched) were obtained from the Koch Institute Preclinical Core and used for all studies. A549 cells were cultured under standard conditions and confirmed to be mycoplasma-free prior to injection. Cells were harvested at 70-80% confluence and resuspended in a 1:1 mixture of 1X PBS and growth factor-reduced Matrigel (Corning, 356231) at a final concentration of 2 × 10^7^ cells/mL. A total of 100 µL of cell suspension was injected subcutaneously into each flank of each mouse using a 27-gauge needle. 4 mice were injected with A549 expressing H3.3-WT and 5 mice were injected with A549 expressing H3.3-Q5H. Tumor volume was monitored every three days using caliper measurements until experimental endpoints were reached, at which point tumors were harvested and weighed. Harvested tumors were processed for immuno-histochemical staining at the Koch Institute Histology Core and analyzed using QuPath software. In parallel, tumor tissue was mechanically dissociated by Dounce homogenization, lysed, and RNA was extracted using phenol-chloroform purification.

### Histology and Immunohistochemistry (IHC)

Tissues were fixed in PBS with 4% (v/v) formaldehyde and then transferred into 70% (v/v) ethanol solution. Tissues were processed, embedded in paraffin and sectioned at 4 µm. Following deparaffinization, tissues were stained with Hematoxylin and Eosin or further processed for IHC using anti-HA tag (CST, C29F4), anti-H3K4me3 (CST, C42D8), and anti-H3K27me3 (CST, C36B11). Images were scanned at 20X using Aperio Digital Pathology Slide Scanner (Leica Biosystems) and processed using QuPath v0.5.1.

### Statistical and Correlation Analyses

Generation of plots and statistical analyses were performed using Prism 10 (GraphPad). Error bars represent standard deviation, unless otherwise noted. Student’s *t*-test (unpaired, two-tailed) was used to assess significance between experimental and control groups and to calculate *P*-values. Multiple comparisons were corrected using the Šídák test for tumor volume comparison. *P*-value < 0.05 was considered statistically significant. To calculate *P*-values for differences in average CUT&RUN signal across promoters and genes, a non-parametric two-tailed Wilcoxon test was used.

## AKNOWLEDGEMENTS

This research was supported by NIH/NIGMS R00 Award (R00-GM140265) (Y.M.S.F.), Ludwig Center at MIT (Y.M.S.F.) (F.J.S.R.), NIH/NCI Core Grant (P30-CA014051) (Y.M.S.F.) (F.J.S.R.), AACR Gertrude B. Elion Cancer Research Award (Y.M.S.F.), V Foundation V Scholar Award (V2022-004, Y.M.S.F) (V2022-028, F.J.S.R.), Howard Hughes Medical Institute (Hanna Gray Fellowship) (F.J.S.R.), Koch Institute Frontier Award (Y.M.S.F/F.J.S.R), Sharp Life Sciences Fellowship (Z.Y.), Ludwig Center Graduate Student Fellowship (Z.Y.), Phillip & Ann Sharp Fellowship (Z.Y.), MIT Presidential Fellowship (Z.Y.) (R.L.B.). We would like to acknowledge all the members of the Soto-Feliciano lab, Ondine Atwa (Sánchez-Rivera lab), and Lindsey Guan (Keating lab) for their help and support in this project. We would also like to thank Alexey A. Soshnev (UT San Antonio), Yu-Jui (Ray) Ho (MSKCC), and Benjamin A. Nacev (University of Pittsburgh) for fruitful discussions on this project. This work was completed with assistance from the following core facilities at Koch Institute at MIT: Swanson Biotechnology Center Flow Cytometry Core Facility (Glenn Paradis and Michele Griffin), Preclinical Modeling (Aurora Burds-Connor), Barbara K. Ostrom (1978) Bioinformatics & Computing Facility (Charlie Whittaker), and BioMicro Center (Stuart Levine).

## AUTHOR CONTRIBUTIONS

Z.Y., F.J.S.R., and Y.M.S.F. conceived the study. Z.Y. conducted patient analysis, constructed the histone mutations library, purified recombinant Tn5, conducted and analyzed all screens. Z.Y. and D.L. conducted screen validation experiments. Z.Y., A.K., and C.E.F. conducted and analyzed mouse experiments; Z.Y. and R.L.B. conducted and analyzed RNA-Seq and CUT&RUN experiments; F.B. performed nucleosome stability computational predictions. Z.Y., M.R.B., and A.A.O. cloned plasmids and transduced cell lines. Z.Y., A.K., R.L.B., F.B., and C.E.F. prepared the figures. L.F., L.G., and A.L.P. conducted experiments. Y.M.S.F., F.J.S.R., A.E.K. supervised the study. Z.Y. and Y.M.S.F. wrote the manuscript with input from all the authors.

---

**Figure S1:**
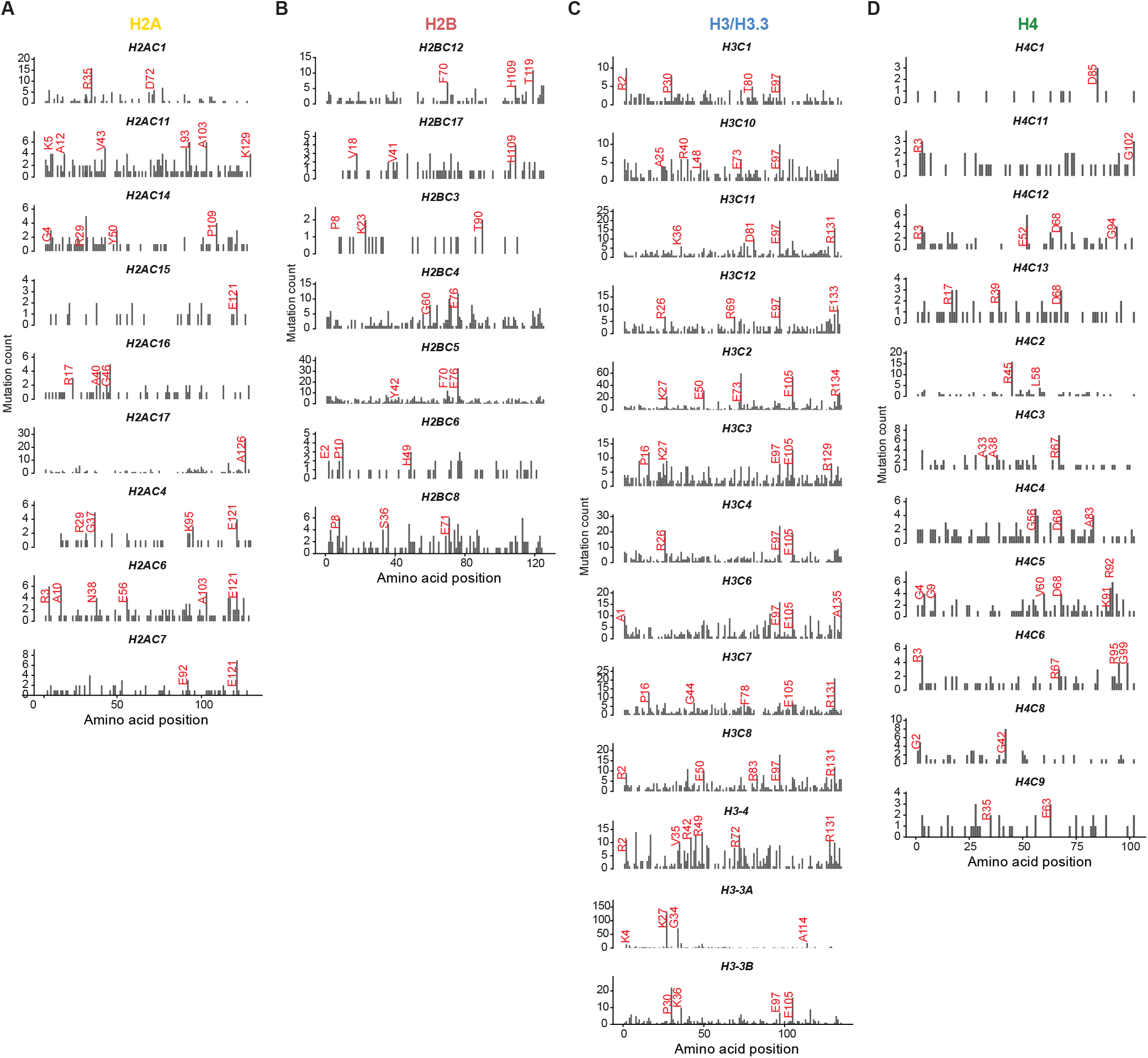
Cancer-associated histone mutations show unique distribution across histone genes. **(A-D)** Mutation profiles for individual histone genes in the H2A **(A)**, H2B **(B)**, H3/H3.3 **(C)**, and H4 **(D)** families. For each gene, the *x*-axis indicates the amino acid position, and the *y*-axis indicates mutation count. Recurrent mutation hotspot residues are labeled in red, highlighting gene-specific patterns of cancer-associated histone mutation across the histone families.

**Figure S2:**
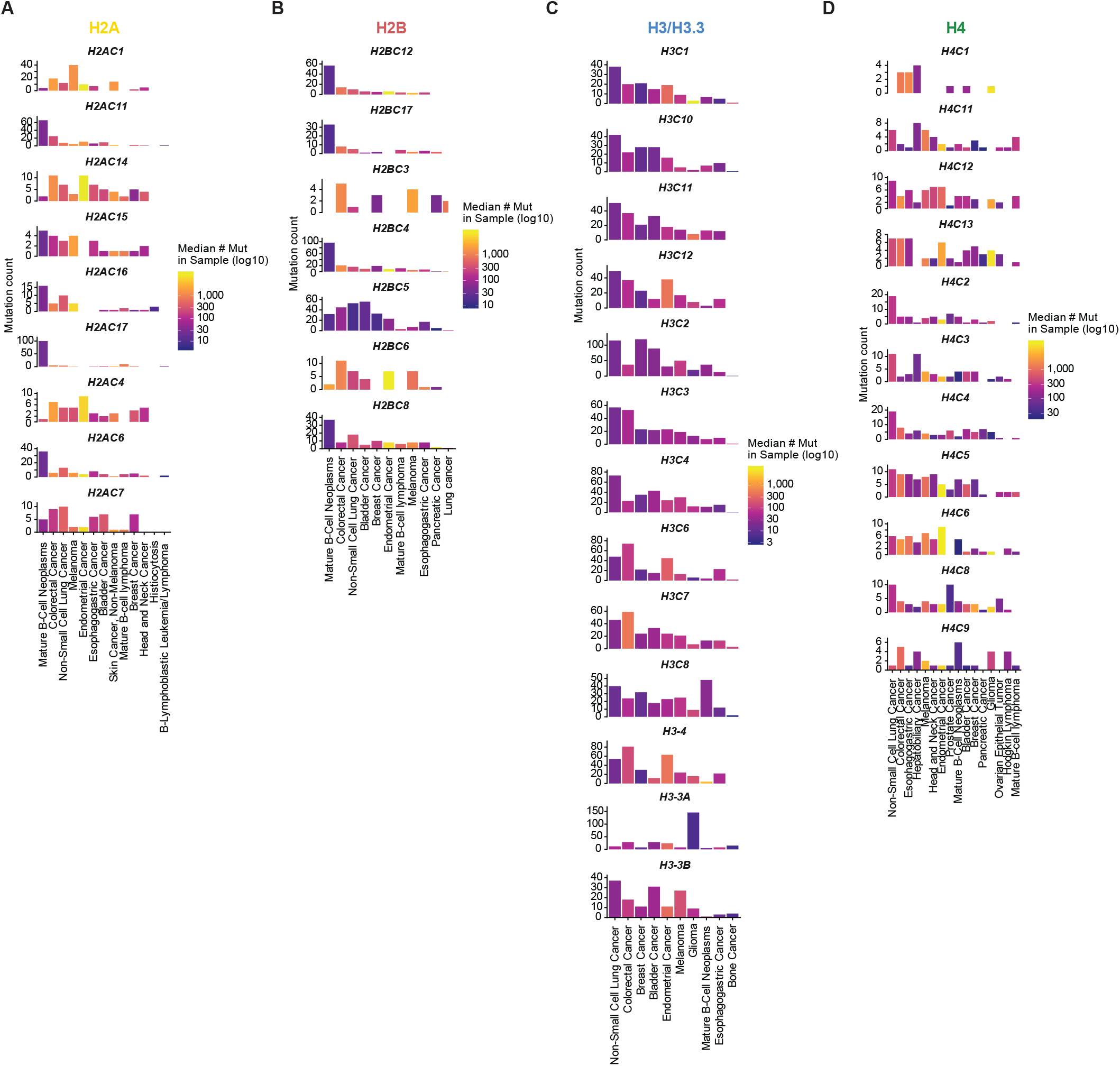
Cancer-associated histone mutations show unique cancer-type enrichment and distinct tumor mutational burden across histone genes. **(A-D)** Horizontal bar plots showing the top five enriched cancer types for each histone gene in the H2A **(A)**, H2B **(B)**, H3/H3.3 **(C)**, and H4 **(D)** families. Bar length represents the mutation count observed for each cancer type-histone gene pair. Bar color indicates the median number of mutations per tumor sample for that cancer type, shown on a log_10_ scale.

**Figure S3:**
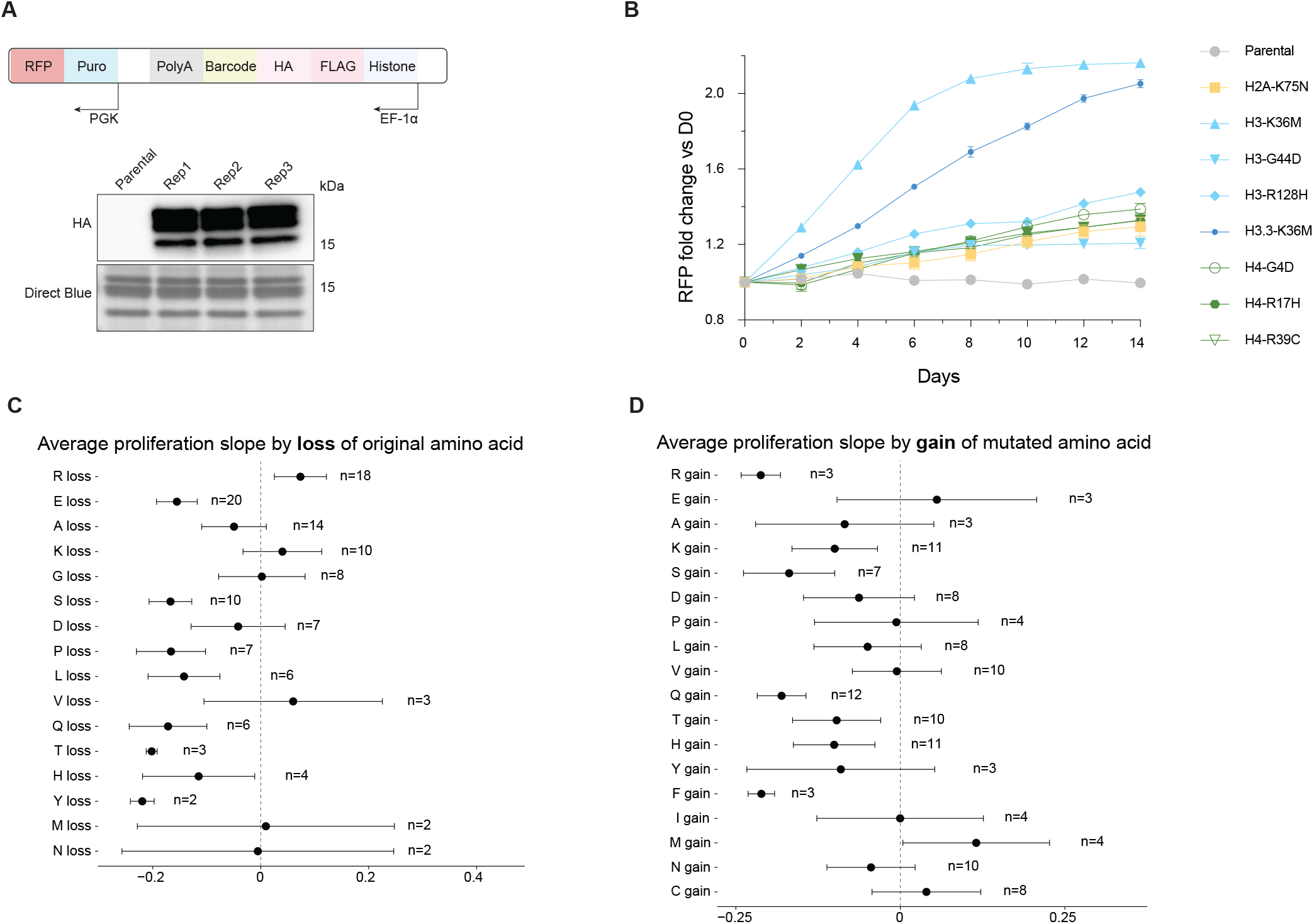
Proliferation screen identifies selective drivers and amino acid substitution patterns associated with proliferation phenotypes. **(A)** Construct design (inverted expression cassette) for the CHANCLA-303 oncohistone library. Acid-extracted histones from mouse mesenchymal stem cells stably expressing the library in three independent biological replicates were immunoblotted using an antibody for the HA tag. **(B)** Growth competition assays in mouse mesenchymal stem cells (MSCs) expressing individual oncohistone proliferative hits. The proportion of RFP+ cells was quantified every 2 days over 14-day period by flow cytometry. **(C)** Average proliferation slope of proliferation screen hits (*P*-value < 0.05) grouped by their original amino acid. **(D)** Average proliferation slope of proliferation screen hits (*P*-value < 0.05) grouped by their mutant amino acid.

**Figure S4:**
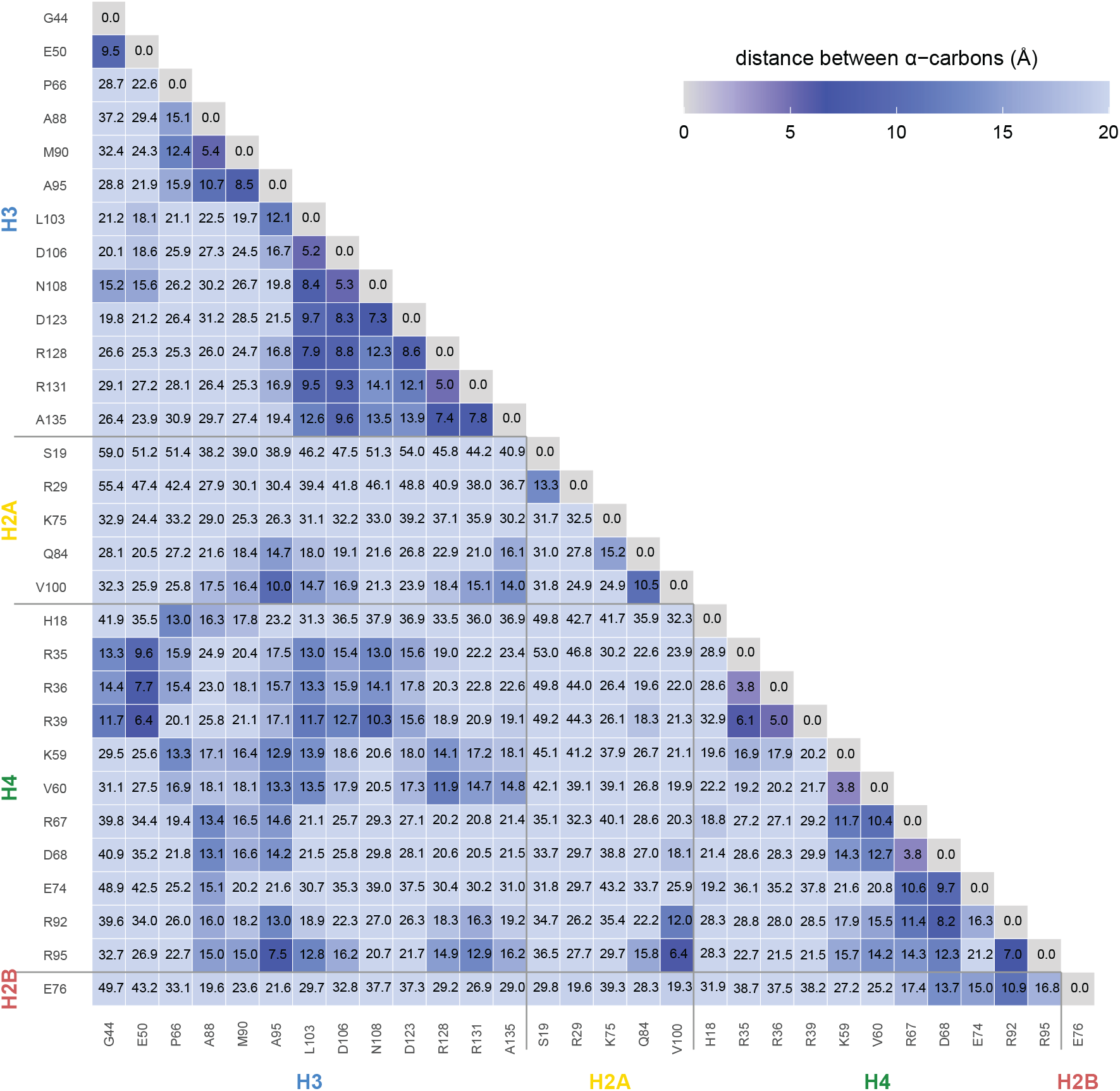
Proximity heatmap showing distances between mutated residues driving proliferation in the nucleosome structure. Spatial proximity heatmap showing distances between proliferative hits in the nucleosome structure (PDB: 2CV5). Plotted residues are shown on the axes and grouped by histone families. Numbers within the grid indicate the calculated distance in angstroms between alpha-carbons (Cα).

**Figure S5:**
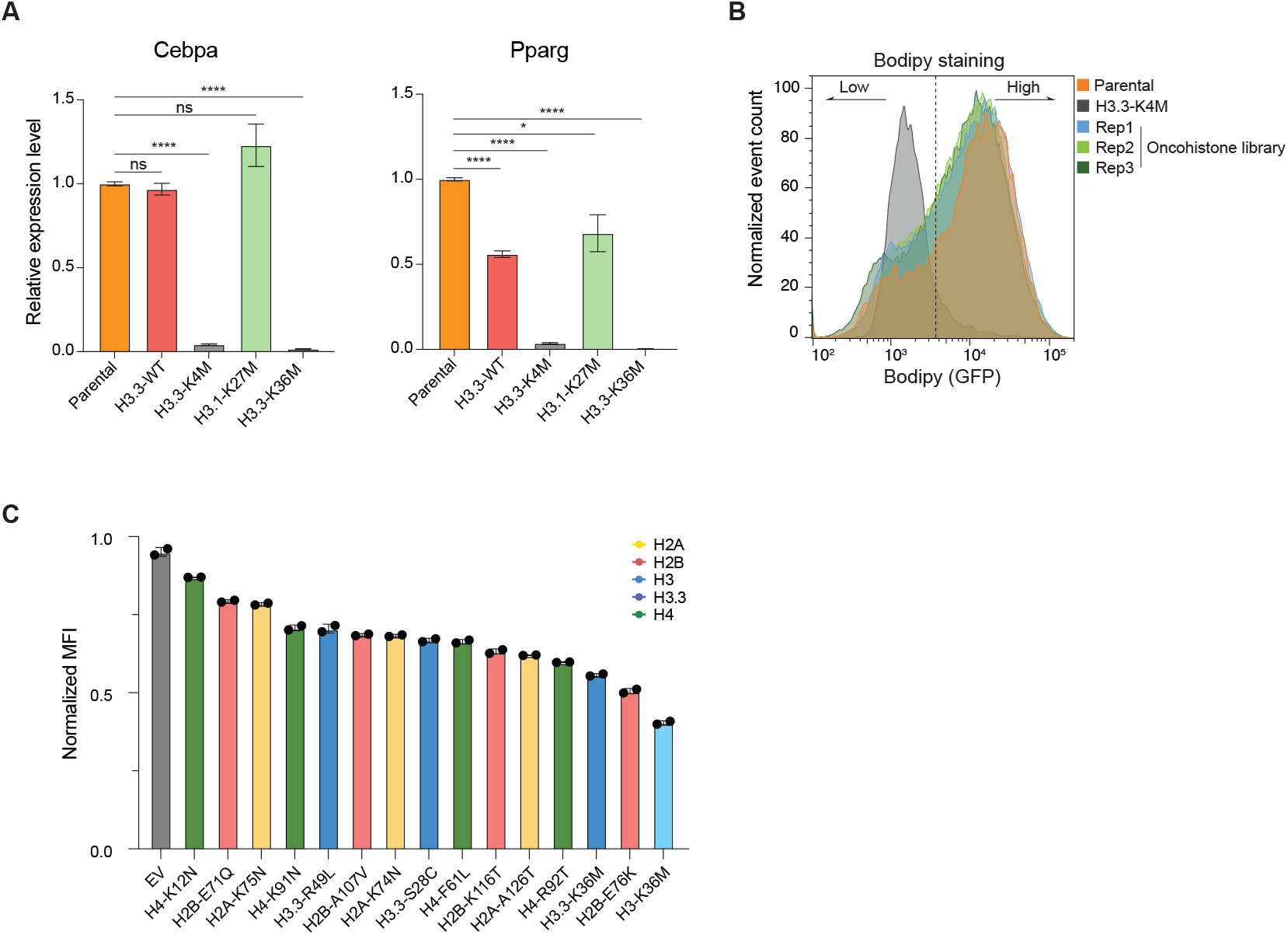
Differentiation screen identifies selective histone mutations impeding adipocyte differentiation. **(A)** qPCR quantification of adipogenesis markers gene expression in mouse mesenchymal stem cells expressing oncohistones, treated with adipocyte differentiation cocktail, and normalized to housekeeping gene *Actb* (n = 3 replicates, SD). **(B)** Normalized event count of mesenchymal stem cells expressing oncohistone library subjected to adipocyte differentiation cocktail, stained with BODIPY, along with parental and *H3*.*3-K4M*-expressing mesenchymal stem cells as gating control for differentiated and poorly differentiated/undifferentiated population sorting, respectively (n = 3 replicates). **(C)** Plot of BODIPY staining of mesenchymal stem cells expressing individual histone mutations subjected to adipocyte differentiation cocktail, MFI measured by flow cytometry and normalized to differentiated parental mesenchymal stem cells.

**Figure S6:**
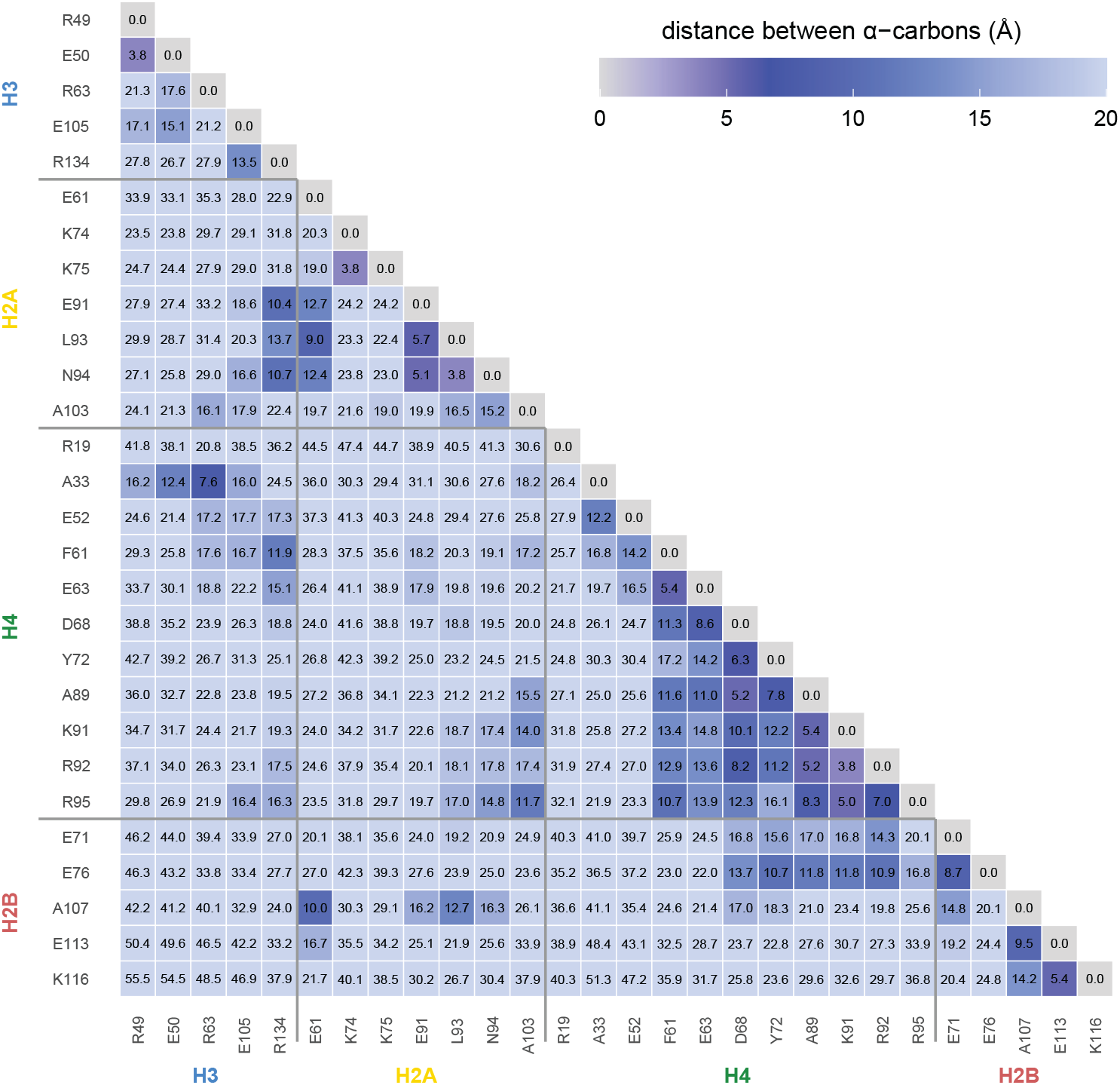
Proximity heatmap showing distances between mutated residues impeding adipocyte differentiation in the nucleosome structure. Spatial proximity heatmap showing distances between differentiation blocker hits in the nucleosome structure (PDB: 2CV5). Plotted residues are shown on the axes and grouped by histone families. Numbers within the grid indicate the calculated distance in angstroms between alphacarbons (Cα).

**Figure S7:**
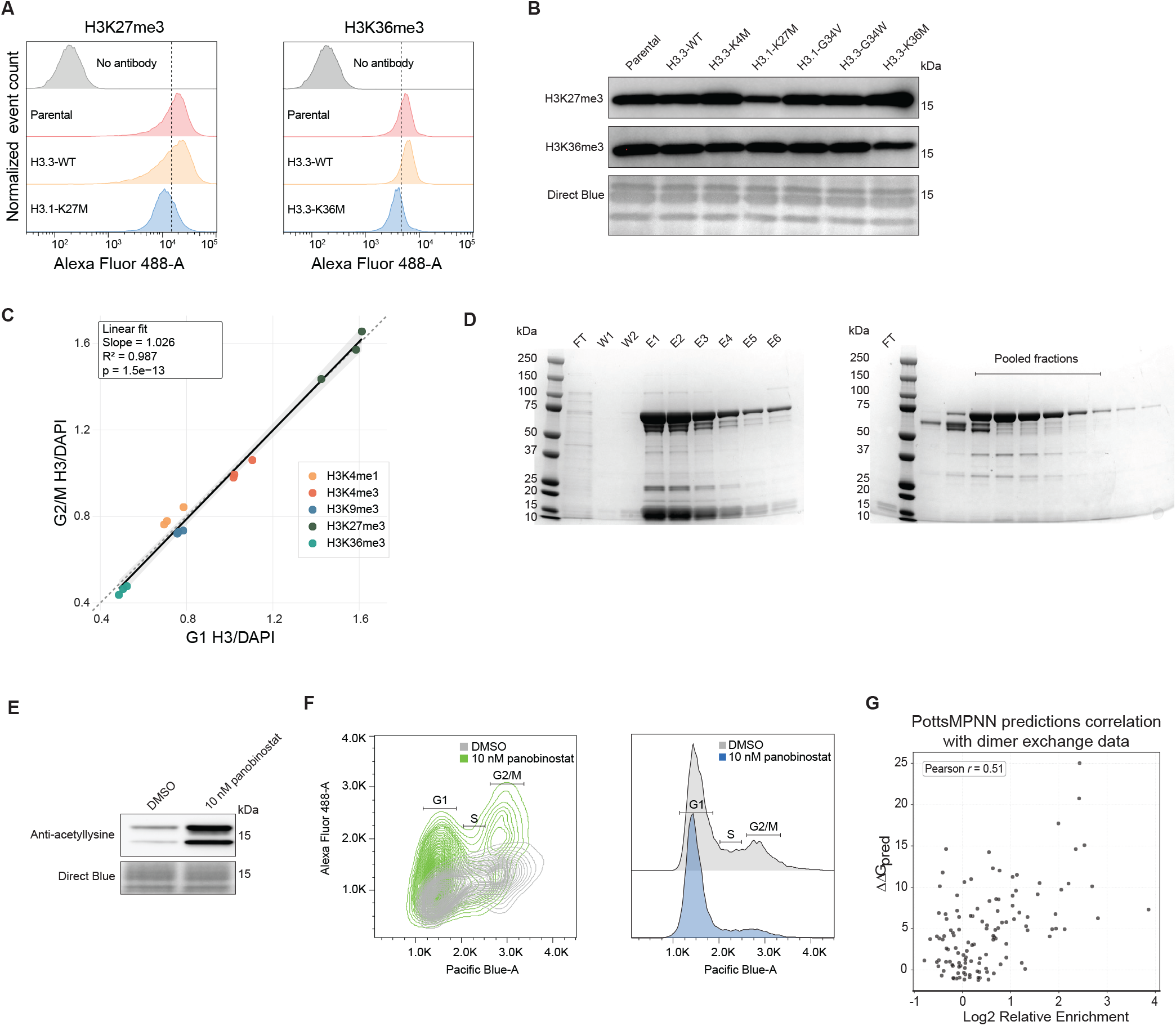
Experimental setup for histone PTM intracellular staining, ATAC-see, and computational modeling of nucleosome stability. **(A)** Flow cytometry analysis of H3K27me3 and H3K36me3 intracellular staining in mesenchymal stem cells expressing mutant histones, *y*-axis showing normalized event count. **(B)** Immunoblot of H3K27me3 and H3K36me3 in acid-extracted histones from mesenchymal stem cells expressing individual mutant histones. **(C)** Linear fit of total histone H3 intracellular staining over DAPI staining in G2/M vs G1 cell population. Individual data points colored by the histone PTM stained in addition to total histone H3. **(D)** Coomassie staining of recombinantly purified Tn5 after Ni-NTA column purification (left) and ion-exchange column purification (right), highlighted with the pooled fractions concentrated for the final product. **(E)** Immunoblot of acetyl-lysine in mesenchymal stem cells treated with DMSO or 10 nM HDACi panobinostat overnight. **(F)** Contour density plot of ATAC-see signal (Alexa Fluor 488) vs DAPI signal (Pacific Blue) in mesenchymal stem cells treated with DMSO or 10 nM HDACi panobinostat overnight. Histogram of DAPI signal (shown right) used to gate cell population into G1, S, and G2/M. **(G)** Correlation plot of PottsMPNN prediction results and published nucleosome dimer exchange data.^12^

**Figure S8:**
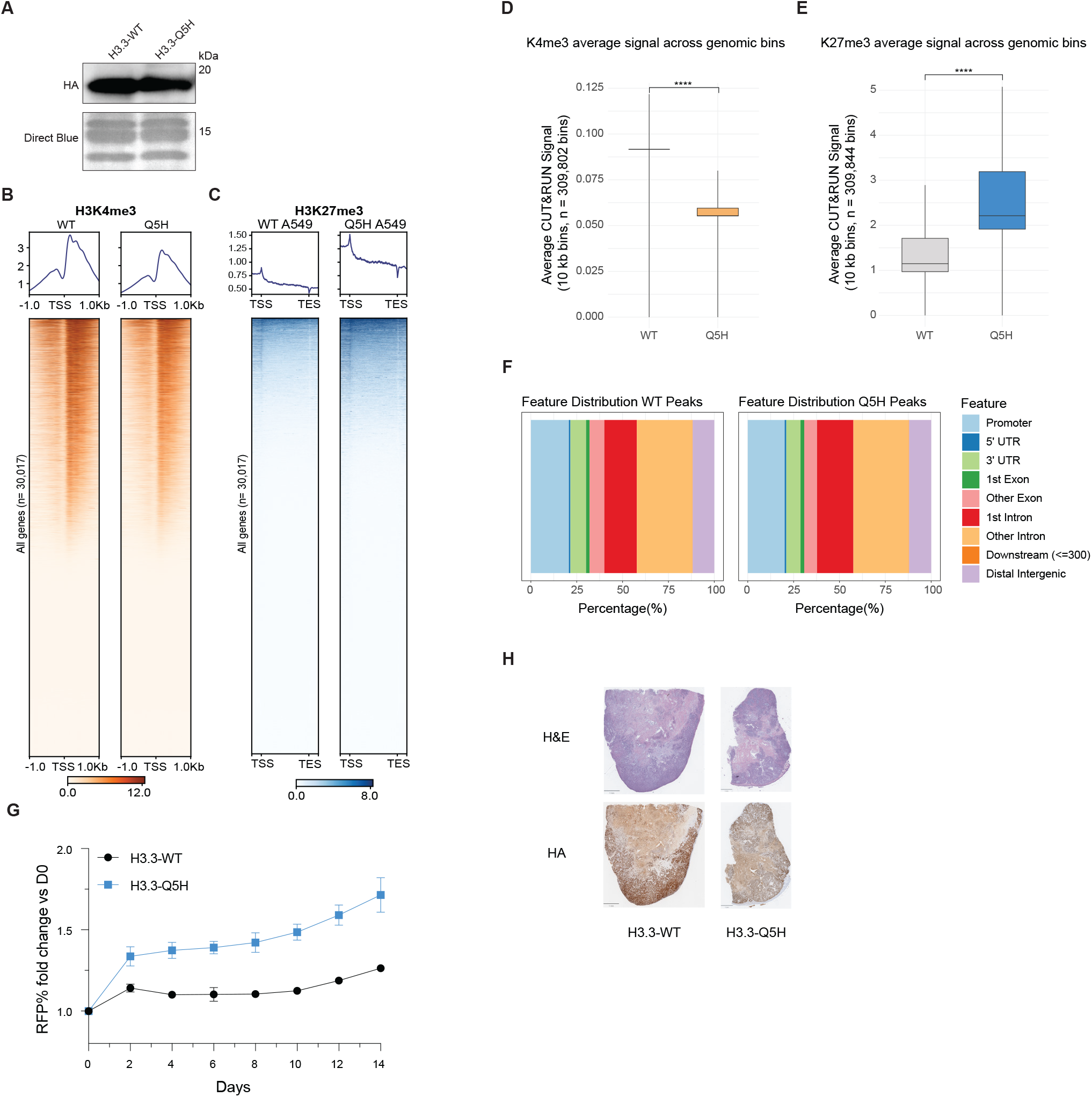
*H3*.*3-Q5H* reshapes global chromatin states in lung adenocarcinoma cells and promotes growth. **(A)** Immunoblot of HA in acid-extracted histones from human A549 cell lines stably expressing *H3*.*3-WT* or *H3*.*3-Q5H*. **(B)** Metagene plots of H3K4me3 CUT&RUN aligned around transcription start sites (TSSs). Heatmap of sorted H3K4me3 CUT&RUN signals at all genes (n = 30,017 genes). **(C)** Metagene plots of H3K27me3 CUT&RUN scaled from TSS to transcription end site (TES). Heatmap of sorted H3K27me3 CUT&RUN signals at all genes (n = 30,017 genes). **(D)** Quantification of H3K4me3 average CUT&RUN signal across all genomic bins (10 kb). **(E)** Quantification of H3K27me3 average CUT&RUN signal across all genomic bins (10 kb). **(F)** Genomic feature distribution plots of H3.3-WT and H3.3-Q5H CUT&RUN peaks. **(G)** *In vitro* growth competition experiments in A549 cell lines expressing *H3*.*3-WT-RFP* or *H3*.*3-Q5H-RFP* (n = 3 replicates, SD). The proportion of RFP+ cells was quantified every 2 days over a 14-day period by flow cytometry. **(H)** H&E staining of representative subcutaneous tumor from A549 cell lines expressing *H3*.*3-WT* or *H3*.*3-Q5H*. IHC staining for HA in the consecutive slide (scale bar represents 1 mm).

